# Cervicovaginal tissue residence imprints a distinct differentiation program upon memory CD8 T cells

**DOI:** 10.1101/769711

**Authors:** Veronica Davé, E. Fabian Cardozo-Ojeda, Florian Mair, Jami Erickson, Amanda S. Woodward-Davis, Amanda Koehne, Andrew Soerens, Julie Czartoski, Candice Teague, Nicole Potchen, Susanne Oberle, Dietmar Zehn, Joshua T. Schiffer, Jennifer M. Lund, Martin Prlic

**Author notes:** Present address: Division of Animal Physiology and Immunology, School of Life Sciences Weihenstephan, Technical University of Munich, 85354 Freising, Germany (Dietmar Zehn).

## Abstract

Tissue-resident memory CD8 T cells (CD8 T_RM_) are critical for maintaining barrier immunity. CD8 T_RM_ have been mainly studied in the skin and gut with recent studies suggesting that the signals that control tissue-residence and phenotype are highly tissue-dependent. We examined the T cell compartment in healthy human cervicovaginal tissue (CVT) and found that most CD8 T cells were granzyme B^+^ and TCF-1^-^. To address if this phenotype is driven by CVT tissue-residence, we used a mouse model to control for environmental factors. Using localized and systemic infection models, we found that CD8 T_RM_ in the mouse CVT gradually acquired a granzyme B^+^, TCF-1^-^ phenotype as seen in human CVT. In contrast to CD8 T_RM_ in the gut, these CD8 T_RM_ were not stably maintained regardless of the initial infection route, which led to reductions in local immunity. Our data show that residence in the CVT is sufficient to progressively shape the size and function of its CD8 T_RM_ compartment.

**Summary:** The tissue-resident memory (T_RM_) CD8 T cell compartment in human and mouse cervicovaginal tissue (CVT) is remarkably similar. The CVT T_RM_ compartment is maintained autonomously and does not reach phenoypical or numerical equilibrium. The numerical decline leads to impaired viral control in a secondary challenge.

## Introduction

After infection or immunization, naive CD8 T cells differentiate into several major populations of memory T cells with distinct functions and trafficking patterns. Tissue-resident memory CD8 T cells (CD8 T_RM_) are a major subset defined by the fact that they do not recirculate via blood and lymph, instead remaining in the tissue sites where they were initially seeded by effector T cells during the primary response (Gebhardt et al., 2009; Masopust et al., 2010; Masopust and Soerens, 2019; Szabo et al., 2019; Wakim et al., 2008). Because of their privileged location, CD8 T_RM_ are uniquely poised to respond to subsequent encounters with their cognate antigen by directly killing infected cells (Casey et al., 2012; Steinbach et al., 2016), proliferating to boost the local CD8 T cell pool (Beura et al., 2018; Park et al., 2018), and activating and recruiting additional immune cells from both resident and circulating populations (Ariotti et al., 2014; Schenkel et al., 2013).

Many factors influence the formation of T_RM_, including the route of priming, exposure to antigen and inflammation, and the type of tissue in which residency is established. For example, the presence of local antigen boosts the size and function of T_RM_ populations within the matched tissue site (Davies et al., 2017; Gebhardt et al., 2009; Khan et al., 2016; Schenkel et al., 2013). Similarly, local inflammation enhances the number and function of T_RM_ in an antigen-independent manner (Mackay et al., 2012; Shin and Iwasaki, 2012) and cytokine cues such as TGF-β, IL-15, and CCL5 appear to directly regulate T_RM_ formation and maintenance (Bergsbaken et al., 2017; Iijima and Iwasaki, 2014; Mackay et al., 2015; Zhang and Bevan, 2013).

Despite the importance of local exposure to antigen and inflammation in T_RM_ development, strong evidence also exists that T_RM_ can broadly seed distal tissues after both systemic and local immunization or infection. Systemic immunization and infection models are routinely used in mice to generate T_RM_ populations across many body sites (Masopust et al., 2001; Milner et al., 2017; Pope et al., 2001; Reinhardt et al., 2001; Steinert et al., 2015). Likewise, localized infections such as herpes simplex virus (HSV) and vaccinia virus seed CD8 T_RM_ at distal tissue sites, albeit to a much lower extent than at the primary site of infection (Davies et al., 2017; Jiang et al., 2012; Khan et al., 2016; Masopust et al., 2004). Together, these studies suggest that T_RM_ can develop within diverse tissue sites across a range of levels of inflammation and antigen availability, and it is currently unclear how the combination of these variables may ultimately tune the characteristics of the resulting T_RM_ population.

The tissue of residence itself also impacts the phenotype of CD8 T_RM_ populations. CD8 T_RM_ have been extensively studied in skin of both humans and mice, and it is now well-known that skin-resident CD8 T cells express the putative residency markers CD69 and integrin α_E_ (CD103), respond to local HSV infection, and rely on TGF-β and IL-15 signals for their development (Gebhardt et al., 2009; Mackay et al., 2013; Watanabe et al., 2015; Zhu et al., 2013). Many reports suggest that CD8 T_RM_ in other organs may differ in their phenotype or developmental requirements. For example, it is well-documented that CD69 and CD103 are not necessarily expressed by T_RM_ in the uterus or pancreas (Steinert et al., 2015) but may be required by T_RM_ in other organs such as the salivary gland, lung, or kidney (Hofmann and Pircher, 2011; Takamura et al., 2016; Walsh et al., 2019). Likewise, some organs such as the small intestine maintain T_RM_ that constitutively express granzyme B in the absence of antigen or re-challenge (Casey et al., 2012; Kim et al., 1997), a phenotype which has not been observed as robustly in other tissue sites, including the lung and uterus (Beura et al., 2019; Piet et al., 2011). It remains an open question whether CD8 T_RM_ phenotype is inextricably linked to tissue of residence, or whether alterations to the priming method or the inflammatory milieu within a tissue could elicit features such as constitutive granzyme B expression in tissue sites where they are not normally observed.

Mucosal barrier tissues contain large populations of CD8 T_RM_, and pathogens that infect mucosal barrier surfaces represent a significant global health burden. Specifically, the study of CD8 T cell memory within the cervicovaginal tissue (CVT) has broad importance in understanding the pathogenesis and vaccinology of sexually-transmitted infections. Compared to memory CD8 T cells in the upper female reproductive tract and other mucosal and lymphoid tissues, the phenotypic and functional characteristics of memory CD8 T cells in the CVT remain relatively understudied. We recently reported that the CD8 T cell compartment in human CVT includes a subset of CD8 T cells that robustly express granzyme B (Pattacini et al., 2019). It is unclear to what extent the distinctive features we observed in human CVT may be driven by recurring local infections, exposure to inflammatory cues, or a result of signals intrinsically associated with the cervicovaginal microenvironment.

Here, we report that CD8 T cells isolated from healthy human CVT lacked expression of the memory-associated transcription factor TCF-1 and resembled effector or terminally-differentiated cells rather than a self-renewing memory T cell population. To determine if this characteristic was driven by residence in the tissue itself or was a consequence of local exposure to antigenic insults or inflammation in human CVT, we utilized multiple immunization strategies to assess memory CD8 T cell differentiation and maintenance in the mouse CVT. We found that memory CD8 T cells in the mouse CVT at later memory time points closely resembled the CD8 T cells found in human CVT regardless of the initial immunization route and early phenotypic differences. Memory CD8 T cell numbers where stably maintained in the periphery and gut, but gradually declined in the CVT over five months post-immunization, which was associated with a delay in viral control upon HSV vaginal challenge. We conclude that residence in the CVT is sufficient to alter the canonical differentiation and maintenance program of memory CD8 T cells.

## Results

### Healthy human cervicovaginal tissue contains CD8 T cells that express granzyme B and lack TCF-1

In order to study the phenotypic profile of CD8 T cells within human CVT from healthy women without any genital infections, we isolated lymphocytes from human vaginal biopsies via enzymatic digestion (**Fig. S1A**) and observed that the majority of memory T cells in this tissue site expressed the residency markers CD69 and CD103 (**Fig. 1A-B**). A large proportion of these CD8 T cells expressed granzyme B *ex vivo* (**Fig. 1C**) and the majority of granzyme B^+^ cells co-expressed CD103 (**Fig. 1D**), suggesting that they were potentially resident within the tissue rather than transient inflammatory cells. We observed that the memory CD8 T cell compartment in human CVT lacked expression of TCF-1 (**Fig. 1E**), a transcription factor associated with self-renewal potential in CD8 T cells (Zhou et al., 2010). These data indicated that human memory CD8 T cells within the CVT may possess functional and proliferative characteristics distinct from such cells in other tissue locations.

**Figure 1.**
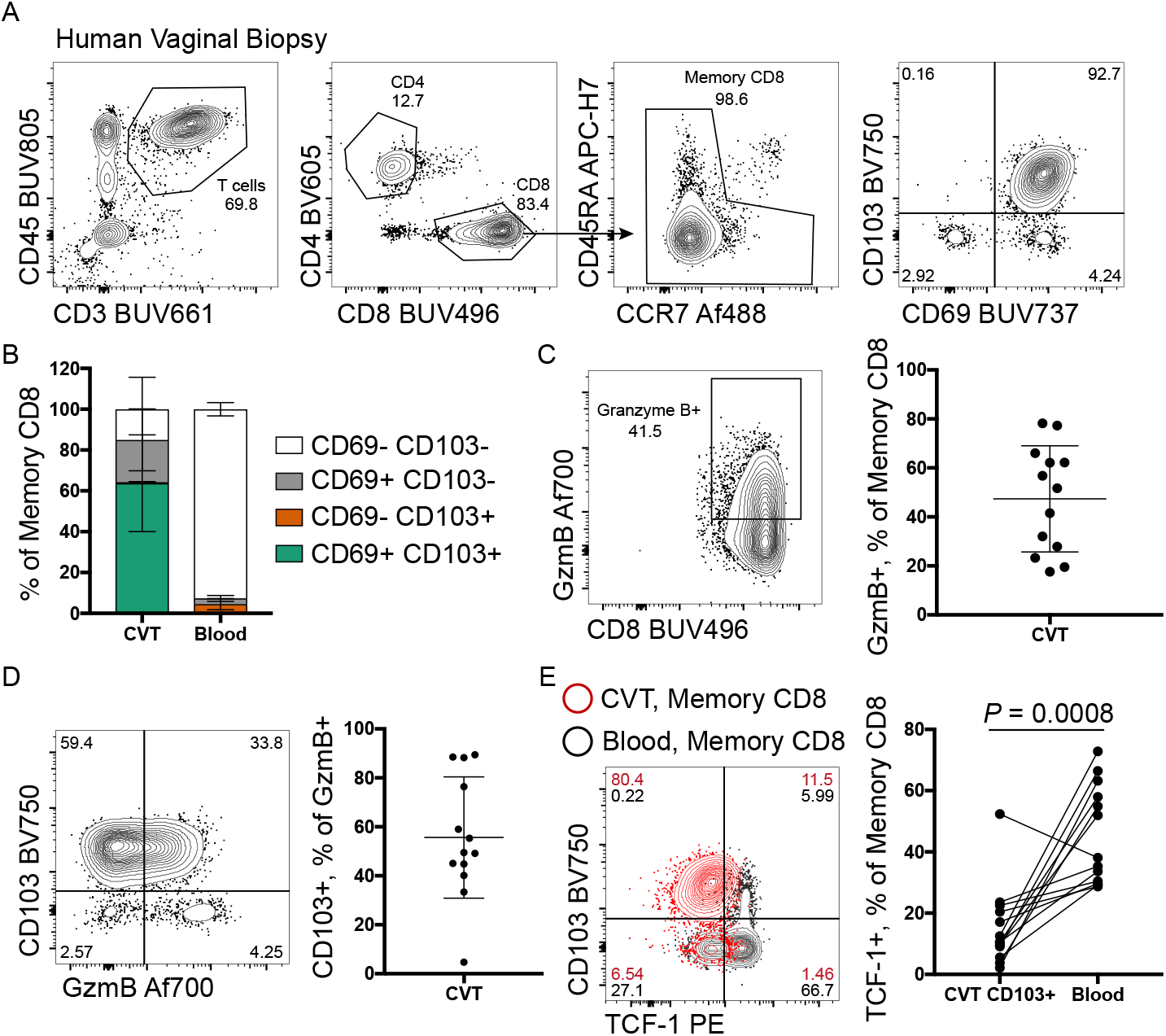
Human cervicovaginal-resident CD8 T cells lack expression of TCF-1. **(A)** Gating strategy to identify memory CD8 T cells in blood and CVT from healthy STI and HSV-2-negative women. **(B)** Relative abundance of CD69 and CD103 subsets in blood and CVT. **(C)** Representative flow plot and abundance of granzyme B^+^ cells within the memory CD8 population in blood and CVT. **(D)** Representative flow plot and abundance of CD103 and granzyme B co-expression in blood and CVT. **(E)** Representative TCF-1 staining and TCF-1^+^ frequency from CD103^+^ subset of CD8 T cells from the CVT compared to memory CD8 T cells in matched blood. Flow plots from A and C-E are from one representative participant. Data from B-E represent pooled results from study participants (n=13). Each dot in C-E represents an individual participant. Error bars represent mean ± SD. *P*-value in E was calculated via paired *t*-test. Exact *P*-values are given for all comparisons.

### Vaginal immunization in mice results in a CD8 T cell compartment that resembles that of human CVT

We next wanted to address if this human CVT CD8 T cell phenotype was attributable to ongoing basal levels of antigenic insult or inflammation in the tissue, or might represent more generalized characteristics of CD8 T cells in CVT. To stringently control for external environmental factors, we sought to generate a comparable population of memory CD8 T cells in mouse CVT. We adoptively transferred naive gBT-I CD8 T cells specific to the SSIEFARL epitope from HSV and primed these cells by vaginal infection with non-lethal thymidine-kinase deficient HSV-2 (HSV-2 TK-) (**Fig. 2A**). One month following infection, HSV gB tetramer^+^ CD8 T cells in the CVT expressed CD69, CD103, and granzyme B in similar proportions to those observed in human CVT (**Fig. 2B-C**). In addition, we found that these CD8 T cells largely lacked expression of TCF-1, especially compared to gB tetramer^+^ cells in the vaginal-draining lymph nodes (dLN) (**Fig. 2D**). Given that these mice had received a vaginal infection, we wondered if this granzyme B^+^ TCF-1^-^ phenotype might be a result of ongoing inflammatory responses in the CVT. Indeed, analysis of hematoxylin and eosin-stained sections of CVT from infected mice revealed that clusters of inflammatory cells remained in the CVT lamina propria as long as 22d after infection (**Fig. 2E**), suggesting that inflammation was not resolved by this time point. Thus, our data suggested that prolonged local tissue inflammation could be driving this distinct memory CD8 T cell phenotype within the CVT.

**Figure 2.**
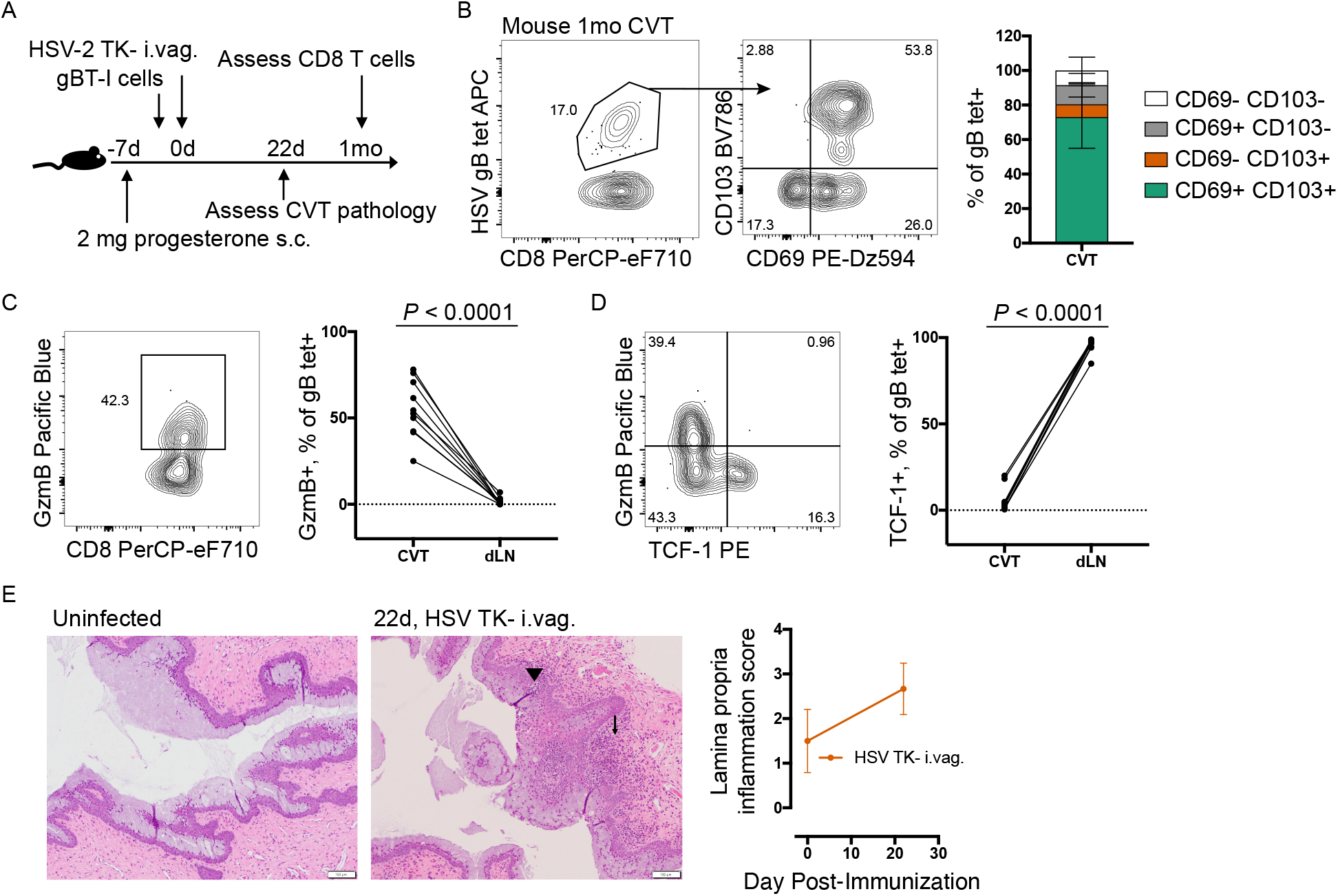
Mouse cervicovaginal-resident CD8 T cells induced by vaginal immunization mirror the phenotype observed in human samples and lack TCF-1. **(A)** Experiment schematic to induce CVT-resident CD8 T cells in mice via vaginal infection with 1.88 × 10^5^ PFU of thymidine kinase-negative HSV-2 186 kpn. **(B)** HSV-specific cells were identified in CVT via staining with an H-2Kb SSIEFARL tetramer conjugated to APC. Left, representative flow plots gated on CD8 T cells. Right, relative abundance of CD69 and CD103 subsets within gB tetramer^+^ CD8 T cells in CVT. **(C)** Representative flow plot and abundance of granzyme B^+^ cells within gB tetramer^+^ CD8 T cells in CVT and CVT-draining lymph nodes (dLN). **(D)** Representative TCF-1 staining and TCF-1^+^ frequency within gB tetramer^+^ CD8 T cells in CVT and dLN. **(E)** Representative images of hematoxylin and eosin-stained sections of CVT at 10X objective and average lamina propria inflammation scores. Arrow indicates coalescing clusters of inflammatory cells with follicular organization. Arrowhead indicates clusters of inflammatory cells within the mucosal epithelium. Flow plots and images from B-E are from one representative mouse. Data from B-D represent pooled results from 2 independent experiments. Each dot in E represents inflammation score averaged from 1-3 mice per time point. Error bars represent mean ± SD. *P*-value in C was calculated via paired *t*-test. Exact *P*-values are given for all comparisons.

### Systemic immunization with LM-gB elicits stable HSV-specific CD8 T cell memory in secondary lymphoid organs and the small intestine, but not the CVT

To determine if prolonged tissue inflammation drives this CVT memory T cell phenotype, we used a systemic immunization approach to prime gBT-I cells in the absence of significant vaginal inflammation. We generated a recombinant strain of *Listeria monocytogenes* that expresses the HSV-derived SSIEFARL peptide (LM-gB) and immunized mice intravenously (**Fig. 3A**). We confirmed that this immunization strategy did not result in detectable levels of vaginal inflammation compared to uninfected mice, and caused significantly less inflammation than observed in mice vaginally immunized with HSV-2 TK- (**Fig. 3B)**. In addition, immunized mice mounted a robust systemic HSV-specific CD8 T cell response, with gB-specific T cells making up an average of 36.4% of CD8 T cells in the blood by 1wk after immunization (**Fig. 3C**). At 1mo after immunization, we could identify these cells in multiple body sites including the CVT and vaginal-draining lymph nodes (**Fig. S1B** and **Fig. 3D**). Again, CD103^+^ gB-specific CD8 T cells in the CVT lacked expression of TCF-1, replicating the phenotype observed in both human vaginal tissue and CVT of vaginally-immunized mice (**Fig. 3E**), despite the lack of tissue inflammation.

**Figure 3.**
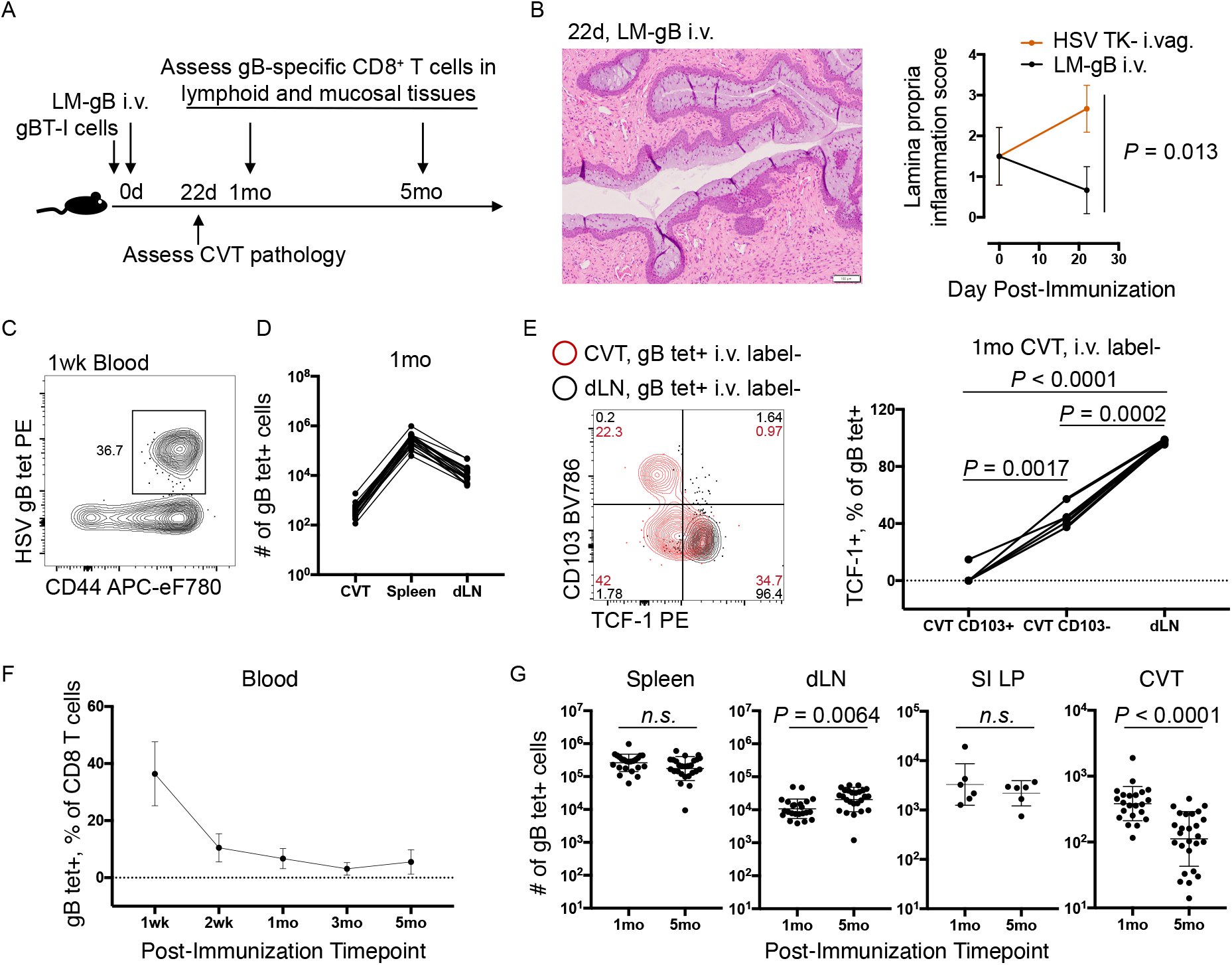
The cervicovaginal CD8 T cell memory compartment of LM-gB-immunized mice is poorly maintained after immunization. **(A)** Schematic of experiment to assess phenotype and maintenance of HSV-specific T cell compartment of mice immunized with 4000 CFU LM-gB i.v. **(B)** Representative image of hematoxylin and eosin-stained sections of CVT at 10X objective and average lamina propria inflammation scores. **(C)** Representative flow plot gated on CD8 T cells from blood collected 1wk after LM-gB immunization. **(D)** gB tetramer^+^ CD8 T cell abundance in spleen, dLN, and CVT 1mo after immunization. **(E)** Representative TCF-1 staining and quantification of TCF-1 expression within CD103^+^ and CD103^−^ subsets of gB tetramer^+^ CD8 T cells in CVT and dLN. **(F-G)** Relative abundance or number of HSV-specific CD8 T cells in blood, spleen, dLN, small intestine lamina propria (SI LP), and CVT 1mo and 5mo after LM-gB immunization. Images and flow plots from B, C, and E are from one representative mouse. Data in D-G are pooled from 2-5 independent experiments. Each dot in B represents inflammation score averaged from 2-3 mice per time point. Each dot in F represents the mean value of 20 mice from a representative experiment. Each dot in D, E, and G represents an individual mouse. Error bars represent mean ± SD. *P*-value in B was calculated via *t*-test. *P-*values in E were calculated with repeated measures ANOVA with Greenhouse-Geisser correction. *P*-values in G were calculated via *t*-test using log_10_-transformed values. Exact *P*-values are given for all values < 0.05.

Given this evidence that memory CD8 T cells in CVT lack significant expression of TCF-1 regardless of immunization route, we next asked whether this memory compartment was deficient in self-renewal and would undergo gradual decay. While the frequency of memory gB-specific CD8 T cells remained relatively stable in the blood following contraction (**Fig. 3F**), and the number of gB-specific CD8 T cells in the spleen, CVT-draining lymph nodes, and small intestine lamina propria (SI LP) beyond 1mo after immunization was stable, this population underwent a threefold loss in number within the CVT, with some mice having very few gB-specific CD8 T cells remaining in the CVT by 5 months post-immunization (**Fig. 3G**). Interestingly, gB-specific T cells in the SI LP also lacked expression of TCF-1 (**Fig. S2A**), suggesting that memory T cells induced by the same immunization may have different requirements for self-renewal.

### Loss of memory CD8 T cells in the CVT occurs mainly among the CD69^−^ CD103^−^ subset

Given that gB-specific CD8 T cells in the CVT continued to decline in number after T cell contraction had concluded in other tissues, we next examined the expression of canonical markers of tissue residency and effector function. To reliably determine what proportion of the gB-specific CD8 T cells in each organ were located within the tissue, we performed intravascular (i.v.) antibody labeling (Galkina et al., 2005) (**Fig. S1C**). We found that an average of 85.6% of the gB-specific CD8 T cells in the CVT were protected from i.v. labeling by 1mo after immunization, similar to the fraction within the small intestine lamina propria (**Fig. 4A**). The proportion of CVT gB-specific CD8 T cells that were protected from i.v. labeling remained stable across time points, suggesting that the smaller population of T cells remaining at late time points continued to be composed of cells that were mostly located within the tissue parenchyma (**Fig. 4A**).

**Figure 4.**
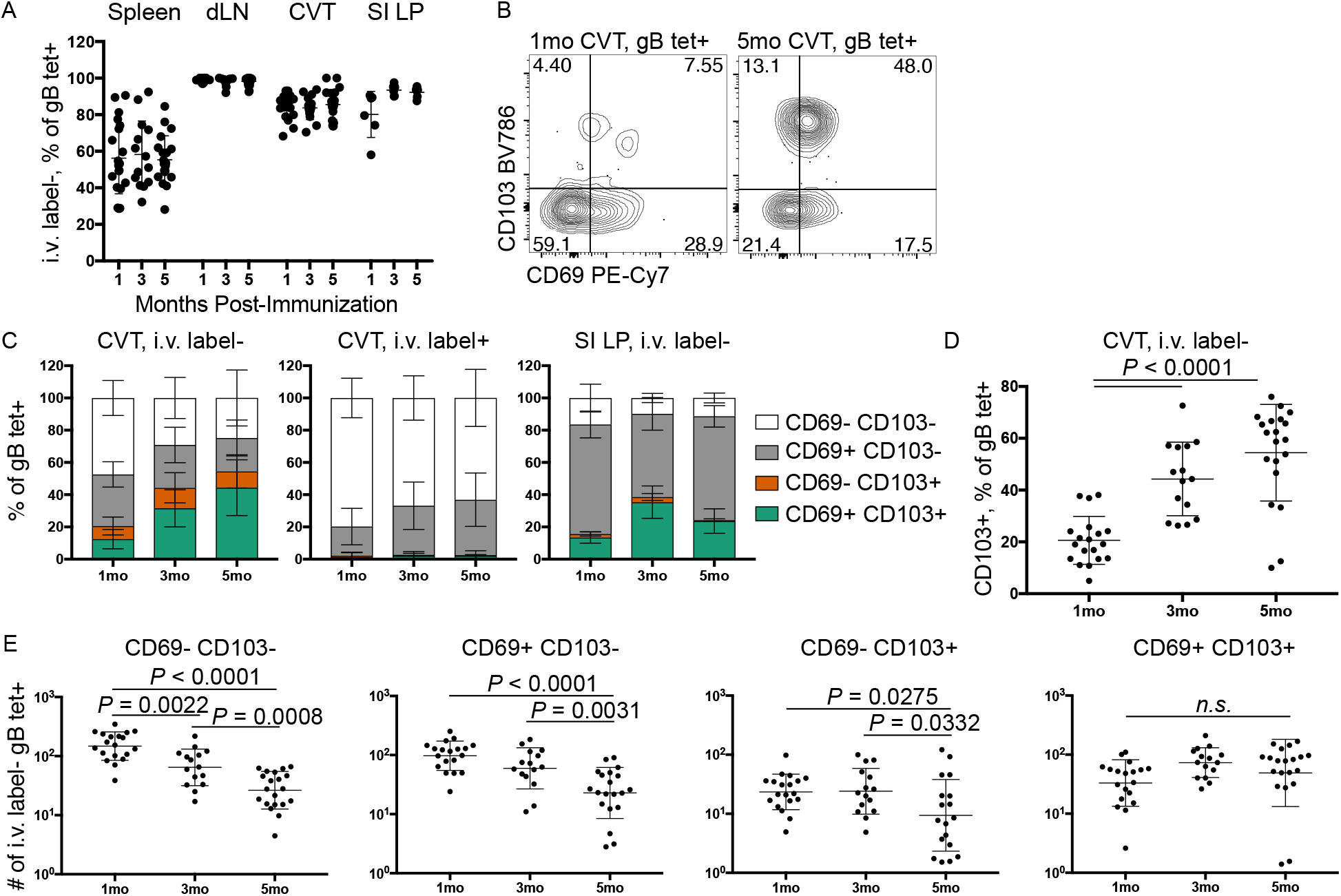
Most HSV-specific CD8 T cells remaining in the CVT by five months after immunization express CD69 and CD103. **(A)** Percentage of i.v. label^−^ HSV-specific CD8 T cells in different lymphoid and non-lymphoid tissues at 1mo, 3mo, and 5mo after immunization based on gating shown in Fig. S1C. **(B)** Representative staining showing CD69 and CD103 expression on HSV-specific CD8 T cells in the CVT 1mo and 5mo after immunization. **(C)** Stacked bar plots showing relative abundance of each of four subsets of HSV-specific CD8 T cells based on CD69 and CD103 expression in the i.v. label^+^ and i.v. label^−^ fractions of the CVT and SI LP. **(D)** CD103 expression on i.v. label^−^ gB tetramer^+^ CD8 T cells in the CVT 1mo, 3mo, and 5mo after immunization. **(E)** Total i.v. label^−^ count of each of four subsets of HSV-specific CD8 T cells based on CD69 and CD103 expression 1mo, 3mo, and 5mo after immunization. Data in A, C, D, and E are pooled from at least 2 independent experiments per time point. Data in B are from one representative experiment. Each dot in A, D, and E represents an individual mouse. Error bars represent mean ± SD. *P*-values in D and E were calculated via ordinary one-way ANOVA with Tukey’s post-hoc test using log_10_-transformed values for numbers and untransformed values for percentages. Exact *P*-values are given for all values <0.05.

We next assessed whether gB-specific CD8 T cells expressed CD69 and CD103. We found that these markers were mostly non-expressed on gB-specific CD8 T cells at the earliest memory time point. However, the average frequency of CD103^+^ cells within the gB-specific CD8 T cell subset increased from 20.6% to 54.4% between 1mo and 5mo after LM-gB immunization (**Fig. 4B-D**). While the i.v.-label^+^ fraction of gB-specific CD8 T cells in the CVT remained almost completely CD103^−^ for the entire duration of the experiment, the shift from a predominantly CD103^−^ population to a predominantly CD103^+^ population was especially clear within the i.v.-label^−^ fraction of CVT gB-specific CD8 T cells (**Fig. 4C-D**) and was mainly attributable to a relative loss of CD103^−^ cells from the CVT over time (**Fig. 4E**). Conversely, the equivalent i.v. label^−^ fraction of gB-specific CD8 T cells in the SI LP, did not undergo a population-level shift towards becoming CD103^+^ (**Fig. 4C**). Finally, the total CD69^+^ CD103^+^ population in the CVT was stably maintained over time (**Fig. 4E**), which could be due to conversion of other subsets to a CD69^+^ CD103^+^ phenotype or the ability to self-renew at a rate sufficient for stable maintenance.

### CVT memory CD8 T cells induced by systemic immunization gradually acquire expression of granzyme B

We next wanted to determine if systemic immunization resulted in constitutive granzyme B expression by CD8 T cells in the CVT, as we observed after vaginal immunization (**Fig. 2**) and as described for tissue-resident CD8 T cells in the SI LP, which express granzyme B in the absence of antigen re-exposure (Casey et al., 2012; Masopust et al., 2006). Few gB-specific CD8 T cells in the CVT expressed granzyme B at 1mo after LM-gB immunization, but the granzyme B^+^ population substantially increased in both number and frequency at later time points (**Fig. 5A-B**). In contrast, a subset of SI LP gB-specific CD8 T cells expressed granzyme B by 1mo after immunization, and expression remained stable thereafter (**Fig. 5A**). As expected, granzyme B expression was rarely observed among gB-specific CD8 T cells in the spleen or dLN, regardless of the memory timepoint (**Fig. 5A**).

**Figure 5.**
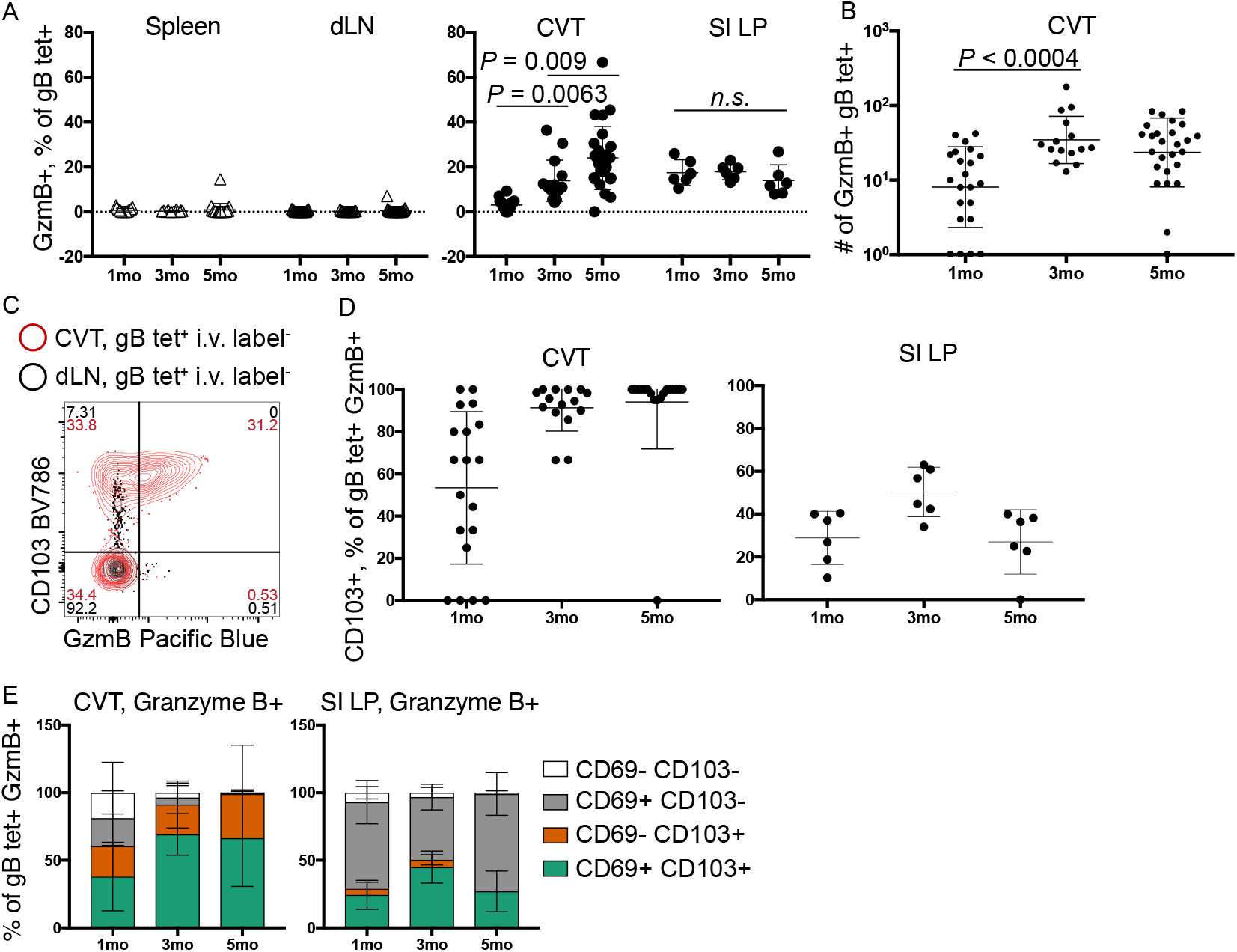
After systemic immunization, cervicovaginal-resident CD8 T cells progressively upregulate granzyme B. **(A)** Frequency of granzyme B^+^ cells among HSV-specific CD8 T cells in spleen, dLN, CVT, and SI LP at 1mo, 3mo, and 5mo post-immunization. **(B)** Number of granzyme B-expressing cells among HSV-specific CD8 T cells at time points after LM-gB immunization. **(C)** Concatenated flow files from 10 mice showing CD103 and granzyme B co-expression among the HSV-specific CD8b i.v. label^−^ populations in the CVT and dLN 5mo after LM-gB immunization. **(D-E)** Frequency of CD69 and CD103 expression within granzyme B^+^ HSV-specific CD8 T cells in the CVT and SI LP 1mo, 3mo, and 5mo after immunization. Data in A-E are pooled from at least 2 independent experiments. Each dot or triangle in A, B, and D represents an individual mouse. Error bars represent mean ± SD. *P*-values in A and B were calculated via ordinary one-way ANOVA with Tukey’s post-hoc test using log10-transformed values for numbers and untransformed values for percentages. Exact *P*-values are given for all values <0.05.

We next examined the phenotypic identity of these granzyme B-expressing cells. By 3mo post-immunization, granzyme B^+^ gB-specific CD8 T cells in the CVT were almost exclusively CD103^+^ (**Fig. 5C-E**), resembling our results in human samples (**Fig. 1D**) and suggesting that these two dynamically-expressed markers were demarcating the same cellular population. By contrast, the granzyme B^+^ population in the SI LP comprised both CD103^+^ and CD103^−^ cells (**Fig. 5D-E**). These data suggest that immunization-induced gB-specific memory CD8 T cells differentiate asynchronously in the CVT relative to the gut mucosa or peripheral lymphoid organs.

### Numeric and phenotypic changes of HSV-specific memory T cells in the CVT lead to delayed protection against viral challenge

Given the decline in the number of gB-specific CD8 T cells in the CVT combined with their shift towards CD103 and granzyme B expression, we next evaluated how these changes affected the immunoprotective response to vaginal HSV-2 challenge. Because LM-gB does not prime HSV-specific CD4 T or B cells responses, this immunization system allowed us to specifically assess the protective effect of CD8 T cells in the absence of other antigen-specific responses. We presumed that the most rapid antiviral responses would likely be mediated by tissue-resident gB-specific CD8 T cells, while circulating gB-specific CD8 T cells would contribute to later responses. We challenged immunized mice at either 1mo (“Early Memory”) or 4mo (“Late Memory”) after immunization (**Fig. 6A**). A similar proportion of mice in each group survived lethal challenge (**Fig. 6B**), demonstrating that gB-specific memory CD8 T cells were sufficient to confer protection against HSV-2 lethality in a subset of immunized mice. However, mice challenged earlier after immunization began to show evidence of faster viral clearance starting as early as 3.5d after challenge (**Fig. 6C**) and underwent greater T cell expansion in the first 2.5d after challenge (**Fig. 6D**). We hypothesized that this was due to quantitative and qualitative differences between the early and late CD8 T cell compartment in the CVT.

**Figure 6.**
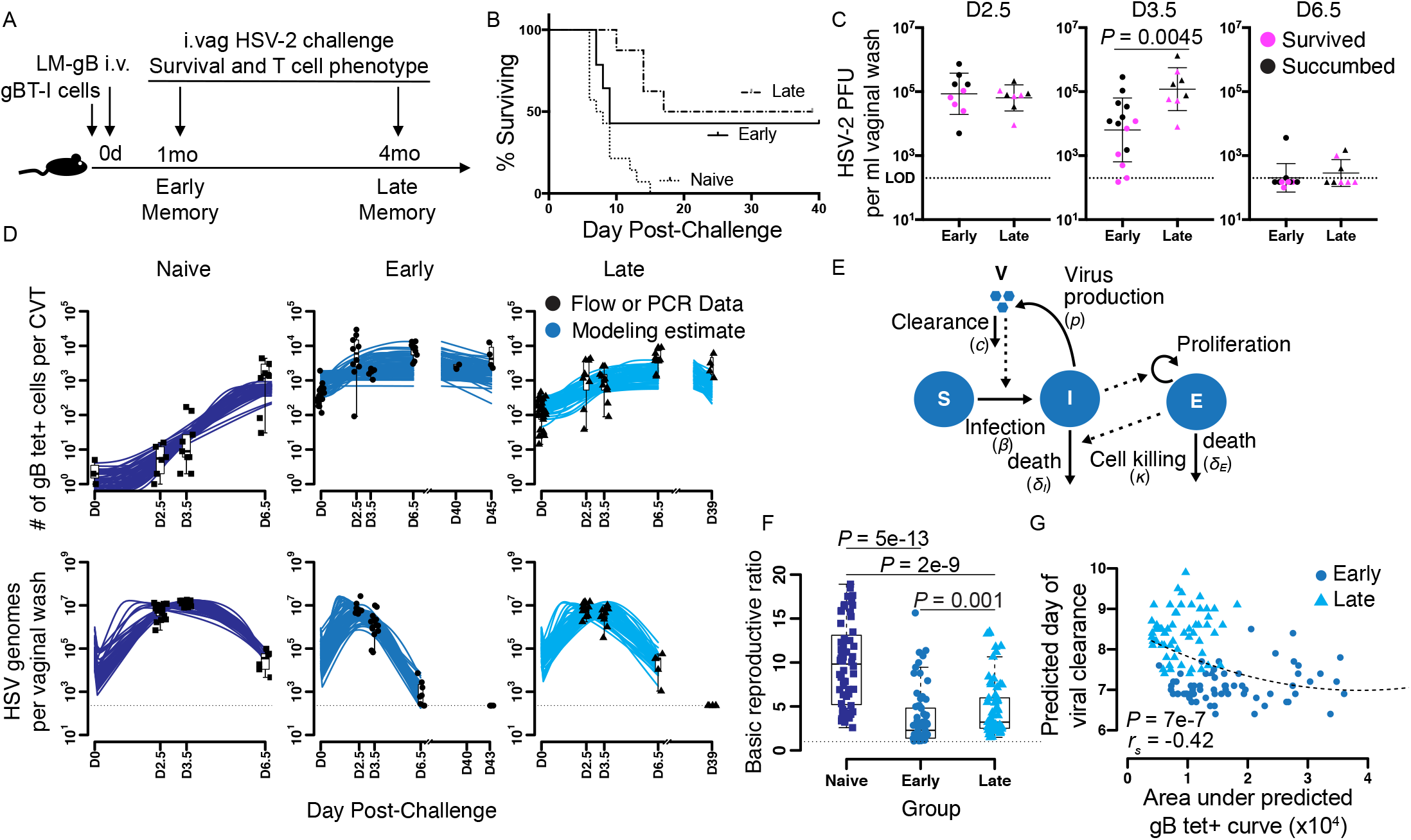
LM-gB immunized mice are protected against severe disease after lethal intravaginal challenge with HSV-2, but the efficacy of the immune response wanes over time. **(A)** Experiment schematic to compare protective efficacy of LM-gB immunization 1mo and 4mo after administration. **(B)** Survival after vaginal HSV-2 challenge of naive and LM-gB immunized mice. **(C)** Vaginal washes were collected after HSV-2 challenge and HSV-2 titer was determined by plaque assay. Pink dots represent mice that survived challenge, as shown in B. **(D)** Number of HSV-specific CD8 T cells (top) and HSV genomes (bottom) in the CVT with modeling estimates overlaid in blue. **(E)** Schematic of mathematical model relating CVT-infiltrating CD8 T cell immune response to HSV-2 viral expansion kinetics. *S* and *I* represent the number of susceptible and infected cells per CVT, respectively; *E*, the number of gB tet^+^ CD8 T cells per CVT; and *V* the number of HSV genomes per vaginal wash. **(F)** Model estimates of basic reproductive ratios (number of new cells infected by one infected cell when introduced into a pool of susceptible cells) for each group of mice. **(G)** Scatter plot of areas under curves (AUC) from D and corresponding predicted day of viral clearance. Dotted line represents a fitted line from quadratic regression. Data in B, C, and D are pooled from at least 2 independent experiments (n=8-14 per group in B). Each dot, square, or triangle in B, C, and D represents data from an individual mouse. Each dot, square, or triangle in F and G represents one model estimate. Error bars in C represent mean ± SD. *P*-values in C were calculated via unpaired *t*-test using log_10_-transformed values. *P*-values in F were calculated via pairwise-corrected Mann-Whitney’s test using Bonferroni’s correction. *P*- and *r*_*s*_-value in G were calculated via Spearman’s rank correlation. Exact *P*-values are given for all values <0.05.

To better understand the relationship between CVT gB-specific CD8 T cells and viral control, we built upon our previously described approaches to model the acute T cell response to HSV-2 infection (Schiffer et al., 2010; Schiffer et al., 2013) and used the mathematical model in **Equation 1** to characterize the different gB-specific T cell response kinetics between the early and late memory groups. In this model, susceptible cells in the CVT are infected by free HSV-2, allowing them to produce new virus or be killed by HSV-specific CD8 T cells (**Fig. 6E**). We performed 100 model-fit rounds to all observations simultaneously for each group for experimentally-determined CVT gB-specific CD8 T cell counts and viral load and overlaid these predictions onto our experimental data (**Fig. 6D)**. In naive mice, gB-specific CD8 T cells were absent from or rare in the CVT at 0-3.5d after challenge but were abundant by 6.5d after challenge (**Fig. 6D**, top left), corresponding with the entry of effector T cells into the CVT following primary infection. In mice challenged at either early or late time points, gB-specific CD8 T cells increased dramatically in number between 0d and 2.5d after HSV-2 infection, and HSV viral titer fell by 6.5 post-challenge (**Fig. 6D**). The best model fits predicted that naive mice had the highest basic reproductive number, meaning that they had the highest number of new cellular infections predicted to derive from a single HSV-infected cell in a susceptible pool (**Fig. 6F**). In addition, the greater areas under fitted gB tet^+^ T cell curves (shown in **Fig. 6D**, top row) were associated with earlier viral clearance (**Fig. 6G**). These data suggest that the numeric and phenotypic changes we observed within the CVT memory T cell compartment led to a decrease in the efficacy of the anti-viral response in the first days after challenge.

### Progressive differentiation by the memory CD8 T cell compartment in the CVT does not require new input of cells from circulation

To determine if the CVT-specific changes in the memory T cell compartment occurred in a tissue-autonomous manner, we depleted Thy1.1^+^ gBT-I cells from the blood and lymphoid tissues of LM-gB immunized mice 1mo after immunization (**Fig. 7A)**, as previously described (Schenkel et al., 2013). This method depleted splenic and circulating gBT-I cells but left the CVT gBT-I compartment intact (**Fig. 7B-C**) and did not have a major impact on protection against vaginal challenge with HSV-2 (**Fig. S3A-B**). By 3mo after depletion, we observed that a similar level of decay had occurred among the CVT gB-specific CD8 T cell population to that seen in undepleted mice between 1mo and 5mo after immunization (**Figs. 3G and 7C**). Similar to our previous results, we observed that this decay occurred mostly among the CD103^−^ subset of T cells (**Fig. 7D-E**) and that the persisting cells had upregulated granzyme B by 3mo after immunization (**Fig. 7F**). Low TCF-1 expression remained stable from 1mo to 3mo after immunization but was reduced by gBT-I depletion, suggesting that the small TCF-1^+^ subset in the CVT of immunized mice was partially but not completely attributable to recirculating gBT-I cells (**Fig. 7G**). These data indicate that the granzyme B^+^ subset of gB-specific cells stems from granzyme B^−^ cells that were present in the tissue 1mo after immunization.

**Figure 7.**
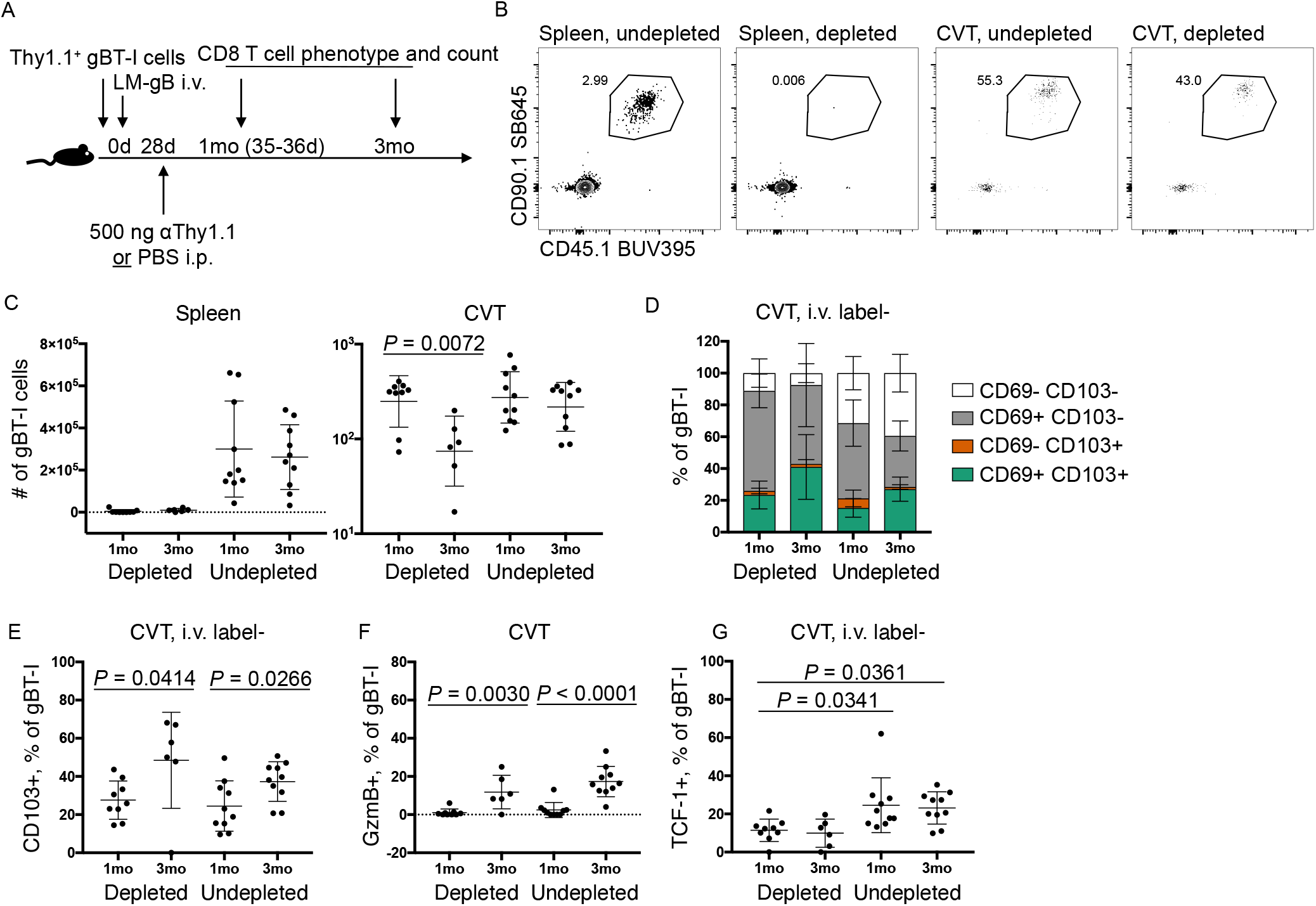
Differentiation and decay of the CVT-resident CD8 T cell compartment is tissue-intrinsic. **(A)** Schematic of experiment to test effect of depleting circulating gBT-I cells. **(B)** Example flow staining to identify gBT-I population within CD8 T cells from spleen and CVT of depleted or undepleted mice 1mo after immunization. **(C)** Number of gBT-I cells recovered from spleen or CVT of mice at indicated time points after immunization. **(D-E)** Relative abundance of CD69 and CD103 subsets within i.v. label^−^ gBT-I cells in CVT at indicated time points. **(F)** Frequency of granzyme B expression among gBT-I cells. **(G)** Frequency of TCF-1 expression among i.v. label^−^ gBT-I cells. Data in C-G are pooled from 2 independent experiments. Flow plots in B are derived from two representative mice. Each dot in C, E, F, and G represents an individual mouse. Error bars represent mean ± SD. *P*-values in C, E, and F were calculated via unpaired *t*-test comparing 1mo to 3mo within each depletion group. *P-*values in G were calculated via ordinary one-way ANOVA with Tukey’s post-hoc test. Exact *P*-values are given for all values <0.05.

### The memory CD8 T cell compartment in the CVT undergoes numeric decay regardless of priming infection and immunization route

Finally, we wished to determine whether site-matched immunization would result in improved maintenance of the CVT memory CD8 T cell compartment. We compared the maintenance of HSV-specific CD8 T cells in the CVT of mice immunized vaginally with HSV-2 TK- or intravenously with LM-gB (**Fig. 8A**). We observed that vaginal immunization with HSV-2 TK- did not result in a larger memory population of HSV-specific CD8 T cells in the CVT than systemic immunization with LM-gB (**Fig. 8B**). In addition, the gB-specific population induced by vaginal immunization was relatively stably maintained in the spleen and SI LP, but underwent decay in the CVT between 1mo and 3mo after immunization (**Fig. 8B**). Despite this decay in the CD8 T cell compartment, HSV-2 TK- immunized mice remained completely protected against lethal vaginal challenge with wildtype HSV-2 (**Fig. S3**), presumably due to CD4 T cells providing substantial protection in this model (Milligan et al., 1998).

**Figure 8.**
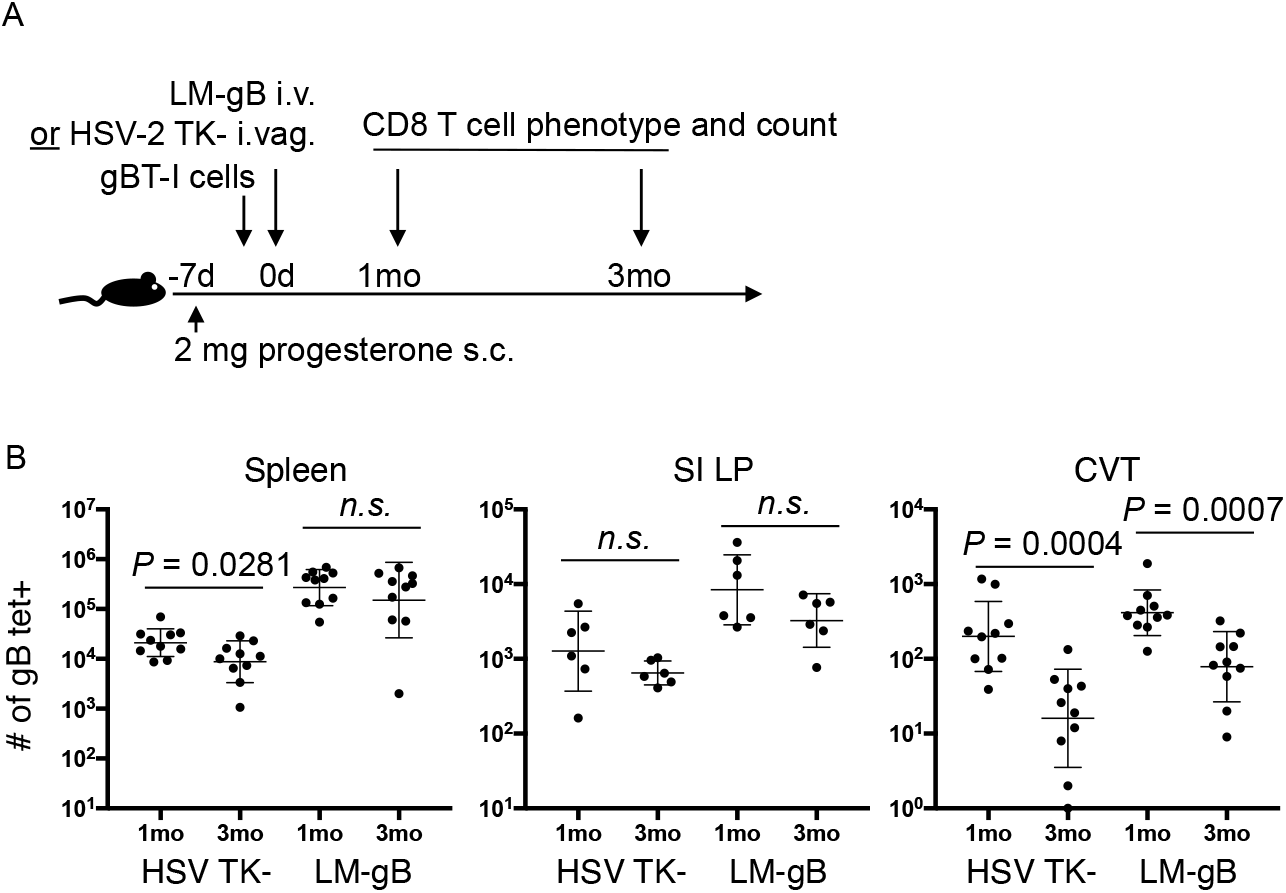
Decay of the CVT-resident CD8 T cell compartment occurs regardless of priming infection and route. **(A)** Schematic of experiment to assess effect of vaginal immunization on longevity of memory CD8 T cells in CVT. **(B)** Number of gB tetramer^+^ cells recovered from spleen, SI LP, or CVT of mice at indicated time points after immunization. Each dot in B represents an individual mouse. Error bars represent mean ± SD. *P*-values in B were calculated via unpaired *t*-test comparing 1mo to 3mo within each immunization group. Exact *P*-values are given for all values <0.05.

## Discussion

Here, we report that CVT microenvironment plays a dynamic role in shaping the numeric, phenotypic, and functional characteristics of the tissue memory CD8 T cells, which thereby impacts their immunoprotective potential. The cervicovaginal mucosa is distinct from other parts of the female reproductive tract and other mucosal tissues in terms of epithelium type and physiology. Specifically, the vagina and ectocervix are composed of type II mucosa and the endocervix and uterus are composed of type I mucosa, with the cervical transformation zone marking the area where the simple columnar epithelial cells of the endocervix meet the stratified squamous epithelial cells of the ectocervix and vagina. In contrast, the gut and the lung, other mucosal tissues commonly studied in the case of CD8 T cell memory, are type I mucosal surfaces (Iwasaki, 2010). In addition to these differences in the structure of the epithelium, there are differences in the complexity and makeup of the microbiota at different mucosal tissue sites (Anahtar et al., 2018), which may also contribute to tissue-specific environmental differences that could affect memory CD8 T cell differentiation and function. Vaginal-resident CD8 T cells in healthy women did not express TCF-1 (**Fig. 1**), which could either simply indicate recent activation of these T cells (recent infection) or a memory population with an impaired ability to self-renew. Importantly, given that vaginal CD8 T cells from multiple donors uniformly lacked TCF-1 expression, it suggested that a tissue-driven phenotype was more likely than a recent infection in a cohort of healthy donors with no current genital infections. Using a mouse model, we similarly found that both vaginal (**Fig. 2**) and systemic (**Fig. 3**) immunization resulted in a TCF-1^low^ phenotype among CD8 T cells in the mouse CVT. These data demonstrate that the mouse and human memory CD8 T cell phenotype are remarkably well conserved in the CVT.

Over the course of 5 months following immunization, antigen-specific memory CD8 T cells in the CVT numerically declined and gradually acquired expression of CD103 and granzyme B. Conversely, HSV-specific CD8 T cells that were circulating (spleen, LN) or resident in the small intestine were numerically and phenotypically stable over the same time course. This decline in CVT T_RM_ memory occurred at a similar rate even when the circulating gBT-I T cells were selectively depleted, suggesting that peripheral T cells do not reseed the CVT under homeostatic conditions (**Fig. 7**). In addition, the CVT CD8 T cell compartment upregulated granzyme B in the absence of new input from the periphery, supporting the occurrence of tissue-intrinsic memory differentiation. Thus, our data show that memory CD8 T cell maintenance and differentiation in the CVT is ultimately directly and uniquely regulated by the tissue environment. A possible explanation for these differences between the vagina and other mucosal tissues such as the gut is that the CVT tissue microenvironment is lacking signals that are important for resident-memory formation and maintenance (Liu et al., 2018). It is noteworthy that we did not observe strong TCF-1 expression in the gut T_RM_ cells either, but these cells were stably maintained over time. This could either suggest that TCF-1 is not required for self-renewal of T_RM_, or the existence of other compensatory mechanisms to maintain the T_RM_ compartment in certain mucosal tissues. TCF-1 activity is thought to be regulated by β-catenin and the Wnt signaling pathway. However, β-catenin does not regulate the phenotype of memory T cells (Driessens et al., 2010) and β-catenin-deficient T cells mount normal recall responses (Prlic and Bevan, 2011). Since β-catenin is sufficient, but not necessary to activate TCF-1 (Tiemessen et al., 2014), a β-catenin-independent pathway may control TCF-1 activity in tertiary tissues with distinct cell fate outcomes compared to β-catenin-dependent activation. A recent study provides some genetic evidence that TCF-1 negatively regulates CD103 expression in the lung tissue (Wu et al., 2020), indicating the need to further study and dissect the role of TCF-1 in controlling differentiation and maintenance of T_RM_ cells.

Understanding the relationship between T_RM_ in mucosal tissues and the periphery is highly relevant, because memory T cell characteristics and frequencies in the blood are used as a benchmark in clinical studies to assess immune responses to vaccines and to establish correlates of protection (Koup et al., 2011; Lewinsohn et al., 2017; Stephenson et al., 2016; Streeck, 2016). Our data indicate that there is a disconnect between memory CD8 T cell frequency, phenotype, and stability in the periphery and the CVT. The numerical and phenotypical changes of the memory CD8 T cell compartment one month vs. 5 months post-immunization ultimately influenced the rapidity of the immune response to vaginal viral challenge (**Fig. 6**). This may be of particular relevance to the success of a vaccine-induced memory T cell response within the CVT, as the host-pathogen events occurring within this early window likely result in the ultimate success or failure of the protective response (Haase, 2010).

Barrier immunity in the CVT remains poorly understood, but is of particular interest, because sexually-transmitted infections (STI) have a significant impact on global health. We currently lack effective vaccines to prevent the majority of these infections, including HIV, HSV, and bacterial STIs. Our data demonstrate that following systemic immunization, animals were partially protected against HSV-2 lethality by gB-specific memory CD8 T cells (**Fig. 6**).

Importantly, this protective effect was observed despite the lack of HSV-specific CD4 T or B cell memory, elements of the immune response that are often considered crucial for protection against HSV-2 (Dudley et al., 2000; Milligan et al., 1998; Morrison et al., 1998; Petro et al., 2016). In line with this observation, results from participants with recurrent genital HSV-2 infection indeed suggest that CD8 T cell activity within lesions is associated with viral control (Peng et al., 2012; Schiffer et al., 2010; Zhu et al., 2007; Zhu et al., 2013). It is possible that a new focus on vaccine strategies that elicit memory CD8 T cells within the genital mucosa, the site of first exposure to STIs, will increase vaccine efficacy. However, successful design of these approaches, not just in regards to adjuvant and immunization route, but also boosting intervals to optimize protection, relies on understanding how memory CD8 T cells are maintained within these genital tract mucosal tissues.

In summary, we demonstrate that the CVT tissue environment controls memory CD8 T cell differentiation and maintenance, which occurs in a tissue autonomous manner and differs across mucosal tissue compartments. The CD8 T_RM_ compartment in the CVT declines over time with a concomitant decrease in the ability to mediate rapid antigen-specific protection.

## Methods

### Mice

C57BL/6J and B6.PL-*Thy1*^*a*^/CyJ mice were purchased from The Jackson Laboratory and maintained in specific pathogen-free conditions at the Fred Hutchinson Cancer Research Center (FHCRC). All mice used in experiments were female and were 8–12 weeks of age. Mice that rejected transferred Thy1.1^+^ gBT-I cells or had an undetectable HSV-2 titer in vaginal wash at 2.5d post-challenge were excluded from experiments. Experiments were approved by the FHCRC Institutional Animal Care and Use Committee.

### Study population

Women recruited for this study (n=13) consented to vaginal biopsy and blood sampling. Eligibility criteria included aged > 18 to < 45, assigned female sex at birth, in good general health, normal PAP smear within the past 3-5 years, not menopausal, Hepatitis C negative and no report of active genital tract irritation or infection. Additionally, participants were screened to be negative for HIV, HSV-2, bacterial vaginosis, chlamydia, gonorrhea, and trichomonas vaginalis. Subjects were not using any type of steroid or medication for treatment of autoimmune disorders. Samples were not collected during menses. Informed written consent was obtained from all participants. The study and procedures were approved by the FHCRC Institutional Review Board.

### Adoptive CD8 T cell transfers

CD44^low^ CD8 T cells were purified via negative selection from spleen and lymph nodes using mouse CD8a^+^ T Cell Isolation kits (Miltenyi) as previously described (Prlic et al., 2001). 5 × 10^4^ cells were injected into recipient B6 mice via i.v. injection into the tail vein.

### Herpes simplex virus type 2 infections and quantification

Mice were injected subcutaneously with 2 mg of medroxyprogesterone acetate injectable suspension (Greenstone). Seven days later, mice were infected intravaginally with 2 × 10^4^ pfu wildtype HSV-2 (strain 186 syn+) or 1.88 × 10^5^ PFU of thymidine kinase-negative HSV-2 186 kpn to the vaginal canal in a 10 ul volume (Parr et al., 1994). After infection with wildtype HSV-2, mice were monitored daily and clinical disease progression was scored as follows: 0, No sign; 1, slight genital erythema; 2, moderate genital erythema and edema; 3, significant genital inflammation with visible lesion; 4, hind leg paralysis or other severe condition requiring euthanasia; 5, moribund or dead. Mice were euthanized if they were moribund or showed signs of severe disease, including hind leg weakness, hind leg paralysis, or hunched posture. Viral titer was determined by plaque assay on Vero cells (ATCC) or by PCR as previously described (Jerome et al., 2002).

### Listeria monocytogenes immunization

A strain of *Listeria monocytogenes* (LM) was generated to recombinantly express and secrete ovalbumin containing the HSV-2 glycoprotein B-derived peptide SSIEFARL according to previously described methods (Lauer et al., 2002; Zehn et al., 2009). Stocks were diluted to 2 × 10^4^ CFU per ml in sterile PBS before immunization. Naive B6 mice received 4,000 CFU i.v. via tail vein injection. Infectious dose was confirmed by plating on brain-heart infusion media plates and enumerating colonies.

### Labeling of T cells in the vasculature

To discriminate CD8 T cells in the vasculature from those located within tissues, we utilized an intravascular labeling technique as previously described (Galkina et al., 2005). Briefly, mice were injected i.v. with 3 ug anti-CD8b antibody conjugated to either FITC or APC (Clone: H35-17.2, eBioscience) in PBS via the tail vein 2-5 min before euthanasia by CO_2_ inhalation.

### Depletion of Thy1.1-expressing T cells

Mice were injected intraperitoneally with 500 ng anti-Thy1.1 antibody (Clone 19E12, Bio X Cell) to deplete Thy1.1-expressing cells from the circulation and peripheral lymphoid tissues.

### Tissue collection and processing

#### Mouse

For pathologic assessment, murine cervicovaginal tracts (consisting of the vagina, ectocervix, and endocervix) were fixed in 10% formalin, embedded in paraffin, sectioned, and stained with hematoxylin and eosin.

To prepare single cell suspensions, cervicovaginal tracts were minced with scissors and then incubated in 2 ml DMEM medium supplemented with 2% FBS, 2 mg/ml collagenase D (Roche), and 1.5 mg/ml DNase I (Roche) for 30 minutes at 37C with gentle rocking. Single cell suspensions were prepared from small intestines by removing the Peyer’s patches and mesentery, incubated in HBSS with 2% FBS, 5 mM EDTA, and 1 mM dithiothreitol for 15 min at 37C with gentle rocking, followed by collagenase treatment as outlined for the CVT.

#### Human

Blood was collected into acid citrate dextrose tubes. Tissues were transported from the clinic in ice cold PBS and immediately processed upon arrival. Biopsy samples were trimmed to 2mm^2^ pieces and digested with collagenase II (700 units/ml, Sigma-Aldrich) and DNAse I (1unit/ml, Sigma-Aldrich) for two subsequent 30 minute-digestions at 37°C as previously described (McKinnon et al., 2014). Single cell suspensions were then washed and filtered before use.

### Pathologic assessment of mouse vaginal tissue

Hematoxylin and eosin-stained sections of mouse CVT were scored based on signs of inflammation within the vaginal lamina propria as follows: 1, Rare neutrophils and other leukocytes present in the lamina propria; 2, Small clusters of neutrophils and other leukocytes present in the lamina propria; 3, Multifocal clusters of leukocytes beginning to coalesce and/or form follicles within the lamina propria; 4, Leukocytes arranged in sheets within the lamina propria. Scoring was performed by a veterinary pathologist who was blinded to group identities.

### Cell Staining for Flow Cytometry and Sorting

Cells were incubated in LIVE/DEAD fixable amine-reactive viability dye (Invitrogen), blocked for Fc binding (Clone 2.4G2 for mice or TruStain (BioLegend) for human), and then stained with tetramers and antibodies.

The following fluorochrome-conjugated antibodies were used to stain mouse cells for flow cytometry: anti-CD8 (Clone 53-6.7, PerCP-eFluor710), anti-CD4 (Clone GK1.5, APC-eFluor780), anti-CD45.1 (Clone A20, Brilliant Ultraviolet 395), anti-CD45.2 (Clone 104, Brilliant Ultraviolet 737), anti-Thy1.1 (Clone HIS51, FITC or SuperBright645), anti-CD44 (Clone IM7, APC or AlexaFluor700), anti-CD69 (Clone H1.2F3, PE-Dazzle594 or PE-Cy7), anti-CD103 (Clone M290, Brilliant Violet 786), anti-granzyme B (Clone GB11, Pacific Blue), and anti-TCF-1 (Clone C63D9, PE). Anti-CD8, anti-CD4, and anti-Thy1.1 were purchased from eBioscience. Anti-CD45.1, anti-CD45.2, anti-CD44, and anti-CD103 were purchased from BD. Anti-CD69 and anti-granzyme B antibodies were purchased from Biolegend. Anti-TCF-1 antibody was purchased from Cell Signaling Technologies.

The following fluorochrome-conjugated antibodies were used to stain human cells for flow cytometry: anti-CD3 (Clone UCHT1, Brilliant Ultraviolet 661), anti-CD45 (Clone HI30, Brilliant Ultraviolet 805), anti-CD8 (Clone RPA-T8, Brilliant Ultraviolet 496), anti-CD4 (Clone RPA-T4, Brilliant Violet 605), anti-CD45RA (Clone HI100, APC-H7), anti-CCR7 (Clone G043H7, AlexaFluor488), anti-granzyme B (Clone GB11, AlexaFluor700), anti-CD103 (Clone Ber-ACT8, Brilliant Violet 750), and anti-TCF-1 (Clone C63D9, PE). Anti-CD3, anti-CD45, anti-CD4, anti-CD8, anti-CD45RA, anti-CD103, and anti-granzyme B were purchased from BD. Anti-CCR7 antibody was purchased from Biolegend. Anti-TCF-1 antibody was purchased from Cell Signaling Technologies.

Intracellular staining was conducted after fixation and membrane permeabilization using Cytofix/Cytoperm Fixation/Permeabilization Kits (BD). If TCF-1 was included in the staining panel, intracellular staining was conducted after nuclear permeabilization using Foxp3/Transcription Factor Staining Kits (eBioscience) according to the manufacturer’s protocol. Samples were acquired on a FACSymphony instrument (BD) or sorted on a FACSAria (BD). To estimate cell counts, 2 × 10^5^ AccuCheck Counting Beads (ThermoFisher) were added to each sample immediately before acquisition.

### Statistical Analysis

Sample means between two unpaired groups were compared by *t*-test. Sample means between three or more unpaired groups were compared by ordinary one-way ANOVA with Tukey’s multiple comparisons test. Sample means between three paired groups were compared by repeated measures ANOVA with Greenhouse-Geisser correction. Paired mean fluorescence intensity or relative frequency data were analyzed via paired *t*-test. All cell count and viral titer data were log_10_-transformed before graphing and analysis. Error bars represent standard deviations. Exact *P*-values are given for all results where *P* < 0.05. Analysis was performed using Prism 7 (GraphPad Software).

### HSV-2 and CD8 T cell Dynamics Modeling

To understand the virus and gB-specific CD8 T cell dynamic differences between the Naive, Early Memory, and Late Memory groups, we used an acute viral infection model that included a cytotoxic effect of the CD8 T cell compartment. In this model, susceptible cells (*S*) are infected at rate *βVS* by free HSV-2 virus (*V*). Productively HSV-2-infected cells (*I*) have a clearance that is intrinsic or mediated by a humoral response with rate *δ*_*I*_*I* or a clearance mediated by HSV-specific CD8 T cells (*E*) with killing rate *kEI*. Free virus is produced at a rate *pI* and cleared at rate *cV*. HSV-specific CD8 T cells are cleared at rate *δ*_*E*_*E* and proliferate in the presence of infected cells at a maximum rate *ωEI*. We also assumed the proliferation of HSV-specific CD8 T cells is constrained when infected and effector cells grow over saturation levels *I*_50_ and *E*_50_, respectively. Under these assumptions the model has the form (see schematics in **Fig. 6D**):

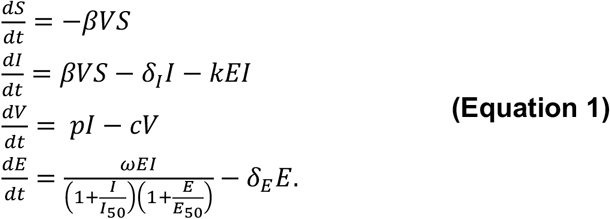

The basic reproductive ratio, *R*_0_, which represents the number of secondary infections produced by an HSV-infected cell when introduced into a susceptible population, was calculated as 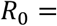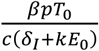. When *R*_0_ < 1, viremia is controlled.

We performed 100 rounds of model fits to observations from all individual animals for each group with different initial parameter guesses in the optimization algorithm. We assumed that all outcomes from the 100 fit rounds were plausible predictions of the data and represented the variability in the infection dynamics in each group. For each round, we re-parameterized the model in function of *R*_0_, and used a nonlinear least-squares approach using the differential evolution and the L-BFGS-B algorithms in R (R Development Core Team) to fit the model and to estimate parameters: *V*_0_, *pT*_0_, *E*_0_, *R*_0_, *δ*, *c*, *k*, *ω*, *δ*_*E*_, *I*_50_ and *E*_50_. Since mice were inoculated with ~2.3×10^6^ HSV genomes, we set that number as the upper limit for *V*_0_. We also constrained the value estimate *E*_0_ to the maximum and minimum HSV-specific CD8 T cell observations at the moment of virus challenge. Finally, we ensured that *R*_0_ > 1 in all fitting rounds. In all simulations *t* = 0 represented the time of the HSV-2 vaginal challenge. From the best fits, we computed the predicted time of viral clearance as the time when viral load crossed the detection limit for the PCR assay of 4615.4 genomes per vaginal wash. We also computed the predicted area under the curve of HSV-specific CD8 T cells using best fits of variable *E* from challenge until the time of viral clearance.

We repeated the fits by exploring different possibilities for constraining HSV-specific CD8 T cell growth. Specifically, we explored a model were CD8 growth is constrained only by a saturation level in HSV-infected cells, i.e. 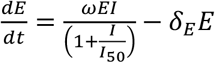, or only by a saturation for the number of effector cells, i.e. 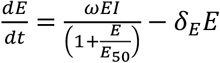. To determine the best and most parsimonious model, we computed the Akaike Information Criteria, 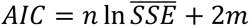, where *m* is the number of parameters estimated, *n* the number of data points in each group and 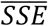 is the average sum of squares error of all the 100 model-fit rounds in each group. We assumed a model had similar support from the data if the difference between its AIC and the best model (lowest) AIC was less than two. We selected and showed results from the model with the lowest AIC.

## Abbreviations

AIC: Akaike information criteria
AUC: area under curve
CVT: cervicovaginal tissue
dLN: CVT-draining lymph nodes
gB: HSV glycoprotein B
HSV-2 TK-: thymidine kinase-deficient herpes simplex virus type 2
LM-gB: *Listeria monocytogenes* strain expressing the SSIEFARL peptide from HSV glycoprotein B
LN: lymph node
SI: small intestine
SI LP: small intestine lamina propria
STI: sexually-transmitted infection

## Acknowledgements

We thank Kristin Weakly for excellent technical assistance and Sean M. Hughes for assistance with statistical analyses. This work was supported by NIH grants R01 AI123323 (MP), R01 AI121129 to (JTS, JML and MP), R01 AI131914 (JML), R01 AI141435 (JML), T32 AI07140 and the Doug and Maggie Walker Fellowship (ASWD), and T32 AI007509 and F31 AI140514 (VD). FM is an ISAC scholar.

## References

Anahtar, M.N., D.B. Gootenberg, C.M. Mitchell, and D.S. Kwon. 2018. Cervicovaginal Microbiota and Reproductive Health: The Virtue of Simplicity. Cell Host Microbe 23:159–168.

Ariotti, S., M.A. Hogenbirk, F.E. Dijkgraaf, L.L. Visser, M.E. Hoekstra, J.Y. Song, H. Jacobs, J.B. Haanen, and T.N. Schumacher. 2014. T cell memory. Skin-resident memory CD8(+) T cells trigger a state of tissue-wide pathogen alert. Science 346:101–105.

Bergsbaken, T., M.J. Bevan, and P.J. Fink. 2017. Local Inflammatory Cues Regulate Differentiation and Persistence of CD8(+) Tissue-Resident Memory T Cells. Cell Rep 19:114–124.

Beura, L.K., N.J. Fares-Frederickson, E.M. Steinert, M.C. Scott, E.A. Thompson, K.A. Fraser, J.M. Schenkel, V. Vezys, and D. Masopust. 2019. CD4(+) resident memory T cells dominate immunosurveillance and orchestrate local recall responses. J Exp Med 216:1214–1229.

Beura, L.K., J.S. Mitchell, E.A. Thompson, J.M. Schenkel, J. Mohammed, S. Wijeyesinghe, R. Fonseca, B.J. Burbach, H.D. Hickman, V. Vezys, B.T. Fife, and D. Masopust. 2018. Intravital mucosal imaging of CD8(+) resident memory T cells shows tissue-autonomous recall responses that amplify secondary memory. Nature immunology 19:173–182.

Casey, K.A., K.A. Fraser, J.M. Schenkel, A. Moran, M.C. Abt, L.K. Beura, P.J. Lucas, D. Artis, E.J. Wherry, K. Hogquist, V. Vezys, and D. Masopust. 2012. Antigen-independent differentiation and maintenance of effector-like resident memory T cells in tissues. Journal of immunology (Baltimore, Md. : 1950) 188:4866–4875.

Davies, B., J.E. Prier, C.M. Jones, T. Gebhardt, F.R. Carbone, and L.K. Mackay. 2017. Cutting Edge: Tissue-Resident Memory T Cells Generated by Multiple Immunizations or Localized Deposition Provide Enhanced Immunity. Journal of immunology (Baltimore, Md. : 1950) 198:2233–2237.

Driessens, G., Y. Zheng, and T.F. Gajewski. 2010. Beta-catenin does not regulate memory T cell phenotype. Nat Med 16:513–514; author reply 514-515.

Dudley, K.L., N. Bourne, and G.N. Milligan. 2000. Immune protection against HSV-2 in B-cell-deficient mice. Virology 270:454–463.

Galkina, E., J. Thatte, V. Dabak, M.B. Williams, K. Ley, and T.J. Braciale. 2005. Preferential migration of effector CD8+ T cells into the interstitium of the normal lung. J Clin Invest 115:3473–3483.

Gebhardt, T., L.M. Wakim, L. Eidsmo, P.C. Reading, W.R. Heath, and F.R. Carbone. 2009. Memory T cells in nonlymphoid tissue that provide enhanced local immunity during infection with herpes simplex virus. Nature immunology 10:524–530.

Haase, A.T. 2010. Targeting early infection to prevent HIV-1 mucosal transmission. Nature 464:217–223.

Hofmann, M., and H. Pircher. 2011. E-cadherin promotes accumulation of a unique memory CD8 T-cell population in murine salivary glands. Proceedings of the National Academy of Sciences of the United States of America 108:16741–16746.

Iijima, N., and A. Iwasaki. 2014. T cell memory. A local macrophage chemokine network sustains protective tissue-resident memory CD4 T cells. Science 346:93–98.

Iwasaki, A. 2010. Antiviral immune responses in the genital tract: clues for vaccines. Nat Rev Immunol 10:699–711.

Jerome, K.R., M.L. Huang, A. Wald, S. Selke, and L. Corey. 2002. Quantitative stability of DNA after extended storage of clinical specimens as determined by real-time PCR. J Clin Microbiol 40:2609–2611.

Jiang, X., R.A. Clark, L. Liu, A.J. Wagers, R.C. Fuhlbrigge, and T.S. Kupper. 2012. Skin infection generates non-migratory memory CD8+ T(RM) cells providing global skin immunity. Nature 483:227–231.

Khan, T.N., J.L. Mooster, A.M. Kilgore, J.F. Osborn, and J.C. Nolz. 2016. Local antigen in nonlymphoid tissue promotes resident memory CD8+ T cell formation during viral infection. J Exp Med 213:951–966.

Kim, S.K., D.S. Reed, W.R. Heath, F. Carbone, and L. Lefrancois. 1997. Activation and migration of CD8 T cells in the intestinal mucosa. Journal of immunology (Baltimore, Md. : 1950) 159:4295–4306.

Koup, R.A., B.S. Graham, and D.C. Douek. 2011. The quest for a T cell-based immune correlate of protection against HIV: a story of trials and errors. Nat Rev Immunol 11:65–70.

Lauer, P., M.Y. Chow, M.J. Loessner, D.A. Portnoy, and R. Calendar. 2002. Construction, characterization, and use of two Listeria monocytogenes site-specific phage integration vectors. Journal of bacteriology 184:4177–4186.

Lewinsohn, D.A., D.M. Lewinsohn, and T.J. Scriba. 2017. Polyfunctional CD4(+) T Cells As Targets for Tuberculosis Vaccination. Front Immunol 8:1262.

Liu, Y., C. Ma, and N. Zhang. 2018. Tissue-Specific Control of Tissue-Resident Memory T Cells. Crit Rev Immunol 38:79–103.

Mackay, L.K., A. Rahimpour, J.Z. Ma, N. Collins, A.T. Stock, M.L. Hafon, J. Vega-Ramos, P. Lauzurica, S.N. Mueller, T. Stefanovic, D.C. Tscharke, W.R. Heath, M. Inouye, F.R. Carbone, and T. Gebhardt. 2013. The developmental pathway for CD103(+)CD8+ tissue-resident memory T cells of skin. Nature immunology 14:1294–1301.

Mackay, L.K., A.T. Stock, J.Z. Ma, C.M. Jones, S.J. Kent, S.N. Mueller, W.R. Heath, F.R. Carbone, and T. Gebhardt. 2012. Long-lived epithelial immunity by tissue-resident memory T (TRM) cells in the absence of persisting local antigen presentation. Proceedings of the National Academy of Sciences of the United States of America 109:7037–7042.

Mackay, L.K., E. Wynne-Jones, D. Freestone, D.G. Pellicci, L.A. Mielke, D.M. Newman, A. Braun, F. Masson, A. Kallies, G.T. Belz, and F.R. Carbone. 2015. T-box Transcription Factors Combine with the Cytokines TGF-beta and IL-15 to Control Tissue-Resident Memory T Cell Fate. Immunity 43:1101–1111.

Masopust, D., D. Choo, V. Vezys, E.J. Wherry, J. Duraiswamy, R. Akondy, J. Wang, K.A. Casey, D.L. Barber, K.S. Kawamura, K.A. Fraser, R.J. Webby, V. Brinkmann, E.C. Butcher, K.A. Newell, and R. Ahmed. 2010. Dynamic T cell migration program provides resident memory within intestinal epithelium. J Exp Med 207:553–564.

Masopust, D., and A.G. Soerens. 2019. Tissue-Resident T Cells and Other Resident Leukocytes. Annu Rev Immunol 37:521–546.

Masopust, D., V. Vezys, A.L. Marzo, and L. Lefrancois. 2001. Preferential localization of effector memory cells in nonlymphoid tissue. Science 291:2413–2417.

Masopust, D., V. Vezys, E.J. Usherwood, L.S. Cauley, S. Olson, A.L. Marzo, R.L. Ward, D.L. Woodland, and L. Lefrancois. 2004. Activated primary and memory CD8 T cells migrate to nonlymphoid tissues regardless of site of activation or tissue of origin. Journal of immunology (Baltimore, Md. : 1950) 172:4875–4882.

Masopust, D., V. Vezys, E.J. Wherry, D.L. Barber, and R. Ahmed. 2006. Cutting edge: gut microenvironment promotes differentiation of a unique memory CD8 T cell population. Journal of immunology (Baltimore, Md. : 1950) 176:2079–2083.

McKinnon, L.R., S.M. Hughes, S.C. De Rosa, J.A. Martinson, J. Plants, K.E. Brady, P.P. Gumbi, D.J. Adams, L. Vojtech, C.G. Galloway, M. Fialkow, G. Lentz, D. Gao, Z. Shu, B. Nyanga, P. Izulla, J. Kimani, S. Kimwaki, A. Bere, Z. Moodie, A.L. Landay, J.A. Passmore, R. Kaul, R.M. Novak, M.J. McElrath, and F. Hladik. 2014. Optimizing viable leukocyte sampling from the female genital tract for clinical trials: an international multi-site study. PLoS One 9:e85675.

Milligan, G.N., D.I. Bernstein, and N. Bourne. 1998. T lymphocytes are required for protection of the vaginal mucosae and sensory ganglia of immune mice against reinfection with herpes simplex virus type 2. Journal of immunology (Baltimore, Md. : 1950) 160:6093–6100.

Milner, J.J., C. Toma, B. Yu, K. Zhang, K. Omilusik, A.T. Phan, D. Wang, A.J. Getzler, T. Nguyen, S. Crotty, W. Wang, M.E. Pipkin, and A.W. Goldrath. 2017. Runx3 programs CD8(+) T cell residency in non-lymphoid tissues and tumours. Nature 552:253–257.

Morrison, L.A., X.J. Da Costa, and D.M. Knipe. 1998. Influence of mucosal and parenteral immunization with a replication-defective mutant of HSV-2 on immune responses and protection from genital challenge. Virology 243:178–187.

Park, S.L., A. Zaid, J.L. Hor, S.N. Christo, J.E. Prier, B. Davies, Y.O. Alexandre, J.L. Gregory, T.A. Russell, T. Gebhardt, F.R. Carbone, D.C. Tscharke, W.R. Heath, S.N. Mueller, and L.K. Mackay. 2018. Local proliferation maintains a stable pool of tissue-resident memory T cells after antiviral recall responses. Nature immunology 19:183–191.

Parr, M.B., L. Kepple, M.R. McDermott, M.D. Drew, J.J. Bozzola, and E.L. Parr. 1994. A mouse model for studies of mucosal immunity to vaginal infection by herpes simplex virus type 2. Laboratory investigation; a journal of technical methods and pathology 70:369–380.

Pattacini, L., A. Woodward Davis, J. Czartoski, F. Mair, S. Presnell, S.M. Hughes, O. Hyrien, G.M. Lentz, A.C. Kirby, M.F. Fialkow, F. Hladik, M. Prlic, and J.M. Lund. 2019. A pro-inflammatory CD8+ T-cell subset patrols the cervicovaginal tract. Mucosal immunology

Peng, T., J. Zhu, K. Phasouk, D.M. Koelle, A. Wald, and L. Corey. 2012. An effector phenotype of CD8+ T cells at the junction epithelium during clinical quiescence of herpes simplex virus 2 infection. J Virol 86:10587–10596.

Petro, C.D., B. Weinrick, N. Khajoueinejad, C. Burn, R. Sellers, W.R. Jacobs, Jr., and B.C. Herold. 2016. HSV-2 DeltagD elicits FcgammaR-effector antibodies that protect against clinical isolates. JCI insight 1:

Piet, B., G.J. de Bree, B.S. Smids-Dierdorp, C.M. van der Loos, E.B. Remmerswaal, J.H. von der Thusen, J.M. van Haarst, J.P. Eerenberg, A. ten Brinke, W. van der Bij, W. Timens, R.A. van Lier, and R.E. Jonkers. 2011. CD8(+) T cells with an intraepithelial phenotype upregulate cytotoxic function upon influenza infection in human lung. J Clin Invest 121:2254–2263.

Pope, C., S.K. Kim, A. Marzo, D. Masopust, K. Williams, J. Jiang, H. Shen, and L. Lefrancois. 2001. Organ-specific regulation of the CD8 T cell response to Listeria monocytogenes infection. Journal of immunology (Baltimore, Md. : 1950) 166:3402–3409.

Prlic, M., and M.J. Bevan. 2011. Cutting edge: beta-catenin is dispensable for T cell effector differentiation, memory formation, and recall responses. Journal of immunology (Baltimore, Md. : 1950) 187:1542–1546.

Prlic, M., B.R. Blazar, A. Khoruts, T. Zell, and S.C. Jameson. 2001. Homeostatic expansion occurs independently of costimulatory signals. Journal of immunology (Baltimore, Md. : 1950) 167:5664–5668.

Reinhardt, R.L., A. Khoruts, R. Merica, T. Zell, and M.K. Jenkins. 2001. Visualizing the generation of memory CD4 T cells in the whole body. Nature 410:101–105.

Schenkel, J.M., K.A. Fraser, V. Vezys, and D. Masopust. 2013. Sensing and alarm function of resident memory CD8(+) T cells. Nature immunology 14:509–513.

Schiffer, J.T., L. Abu-Raddad, K.E. Mark, J. Zhu, S. Selke, D.M. Koelle, A. Wald, and L. Corey. 2010. Mucosal host immune response predicts the severity and duration of herpes simplex virus-2 genital tract shedding episodes. Proceedings of the National Academy of Sciences of the United States of America 107:18973–18978.

Schiffer, J.T., D. Swan, R. Al Sallaq, A. Magaret, C. Johnston, K.E. Mark, S. Selke, N. Ocbamichael, S. Kuntz, J. Zhu, B. Robinson, M.L. Huang, K.R. Jerome, A. Wald, and L. Corey. 2013. Rapid localized spread and immunologic containment define Herpes simplex virus-2 reactivation in the human genital tract. Elife 2:e00288.

Shin, H., and A. Iwasaki. 2012. A vaccine strategy that protects against genital herpes by establishing local memory T cells. Nature 491:463–467.

Steinbach, K., I. Vincenti, M. Kreutzfeldt, N. Page, A. Muschaweckh, I. Wagner, I. Drexler, D. Pinschewer, T. Korn, and D. Merkler. 2016. Brain-resident memory T cells represent an autonomous cytotoxic barrier to viral infection. J Exp Med 213:1571–1587.

Steinert, E.M., J.M. Schenkel, K.A. Fraser, L.K. Beura, L.S. Manlove, B.Z. Igyarto, P.J. Southern, and D. Masopust. 2015. Quantifying Memory CD8 T Cells Reveals Regionalization of Immunosurveillance. Cell 161:737–749.

Stephenson, K.E., H.T. D’Couto, and D.H. Barouch. 2016. New concepts in HIV-1 vaccine development. Curr Opin Immunol 41:39–46.

Streeck, H. 2016. Designing optimal HIV-vaccine T-cell responses. Curr Opin HIV AIDS 11:593–600.

Szabo, P.A., M. Miron, and D.L. Farber. 2019. Location, location, location: Tissue resident memory T cells in mice and humans. Sci Immunol 4:

Takamura, S., H. Yagi, Y. Hakata, C. Motozono, S.R. McMaster, T. Masumoto, M. Fujisawa, T. Chikaishi, J. Komeda, J. Itoh, M. Umemura, A. Kyusai, M. Tomura, T. Nakayama, D.L. Woodland, J.E. Kohlmeier, and M. Miyazawa. 2016. Specific niches for lung-resident memory CD8+ T cells at the site of tissue regeneration enable CD69-independent maintenance. J Exp Med 213:3057–3073.

Tiemessen, M.M., M.R. Baert, L. Kok, M.C. van Eggermond, P.J. van den Elsen, R. Arens, and F.J. Staal. 2014. T Cell factor 1 represses CD8+ effector T cell formation and function. Journal of immunology (Baltimore, Md. : 1950) 193:5480–5487.

Wakim, L.M., J. Waithman, N. van Rooijen, W.R. Heath, and F.R. Carbone. 2008. Dendritic cell-induced memory T cell activation in nonlymphoid tissues. Science 319:198–202.

Walsh, D.A., H. Borges da Silva, L.K. Beura, C. Peng, S.E. Hamilton, D. Masopust, and S.C. Jameson. 2019. The Functional Requirement for CD69 in Establishment of Resident Memory CD8(+) T Cells Varies with Tissue Location. Journal of immunology (Baltimore, Md. : 1950) 203:946–955.

Watanabe, R., A. Gehad, C. Yang, L.L. Scott, J.E. Teague, C. Schlapbach, C.P. Elco, V. Huang, T.R. Matos, T.S. Kupper, and R.A. Clark. 2015. Human skin is protected by four functionally and phenotypically discrete populations of resident and recirculating memory T cells. Sci Transl Med 7:279ra239.

Wu, J., A. Madi, A. Mieg, A. Hotz-Wagenblatt, N. Weisshaar, S. Ma, K. Mohr, T. Schlimbach, M. Hering, H. Borgers, and G. Cui. 2020. T Cell Factor 1 Suppresses CD103+ Lung Tissue-Resident Memory T Cell Development. Cell Rep 31:107484.

Zehn, D., S.Y. Lee, and M.J. Bevan. 2009. Complete but curtailed T-cell response to very low-affinity antigen. Nature 458:211–214.

Zhang, N., and M.J. Bevan. 2013. Transforming growth factor-beta signaling controls the formation and maintenance of gut-resident memory T cells by regulating migration and retention. Immunity 39:687–696.

Zhou, X., S. Yu, D.M. Zhao, J.T. Harty, V.P. Badovinac, and H.H. Xue. 2010. Differentiation and persistence of memory CD8(+) T cells depend on T cell factor 1. Immunity 33:229–240.

Zhu, J., D.M. Koelle, J. Cao, J. Vazquez, M.L. Huang, F. Hladik, A. Wald, and L. Corey. 2007. Virus-specific CD8+ T cells accumulate near sensory nerve endings in genital skin during subclinical HSV-2 reactivation. J Exp Med 204:595–603.

Zhu, J., T. Peng, C. Johnston, K. Phasouk, A.S. Kask, A. Klock, L. Jin, K. Diem, D.M. Koelle, A. Wald, H. Robins, and L. Corey. 2013. Immune surveillance by CD8alphaalpha+ skin-resident T cells in human herpes virus infection. Nature 497:494–497.

