## Supplementary material for "Cervicovaginal tissue residence imprints a distinct differentiation program upon memory CD8 T cells": Suppl Figs 1 -3

**Supplemental Figure 1**

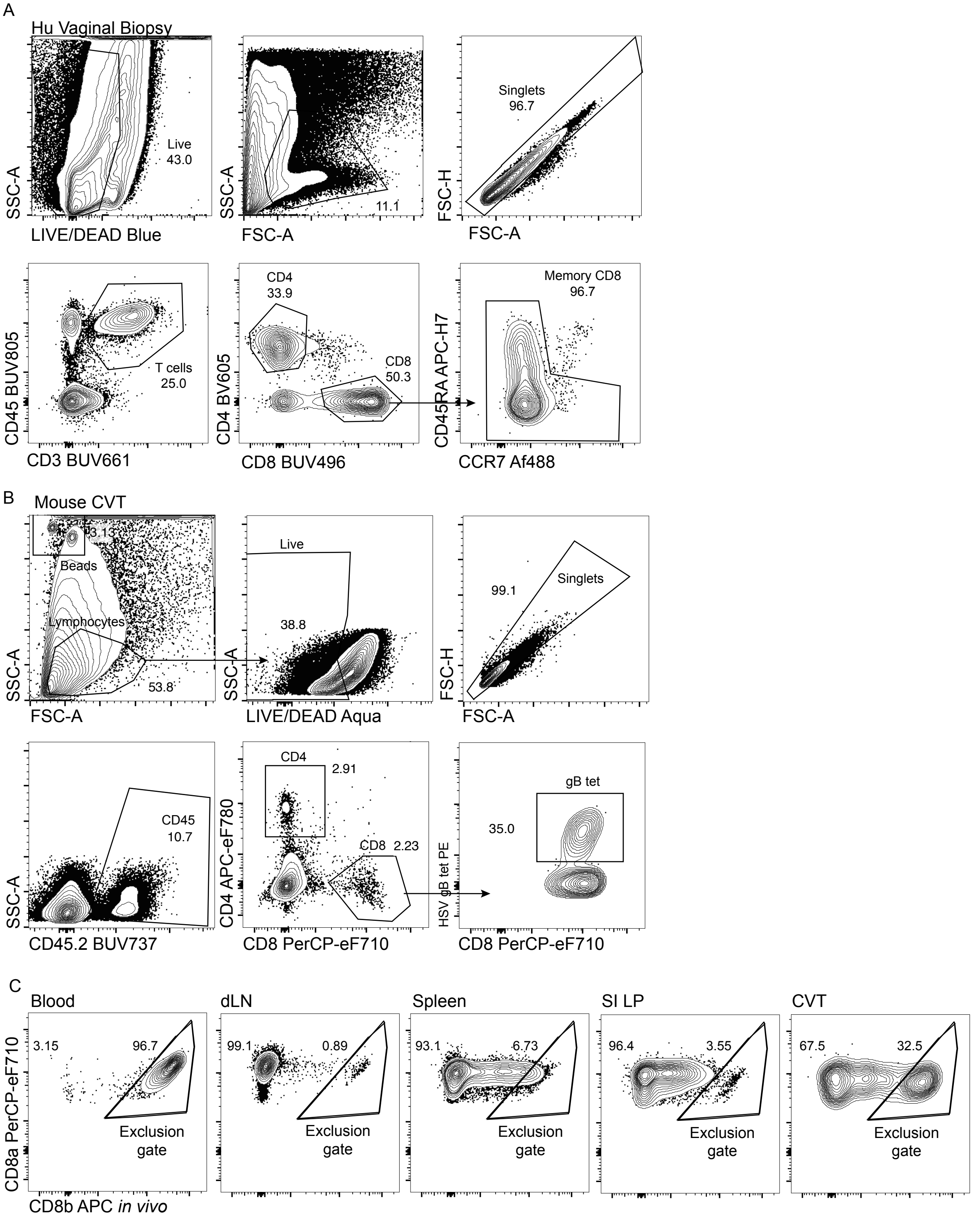


**Figure S1. Example of gating strategies used to identify CD8 T cells in human and mouse CVT.** (A) Representative example of T cell gating strategy from vaginal biopsies from one study participant. (B) Representative example from CVT of mouse sacrificed 1mo after LM-gB immunization. (C) Comparison of intravascular staining using CD8b-APC in blood, dLN, spleen, SI LP, and CVT. Representative staining examples from one mouse sacrificed 5mo after LM-gB immunization.

**Supplemental Figure 2**

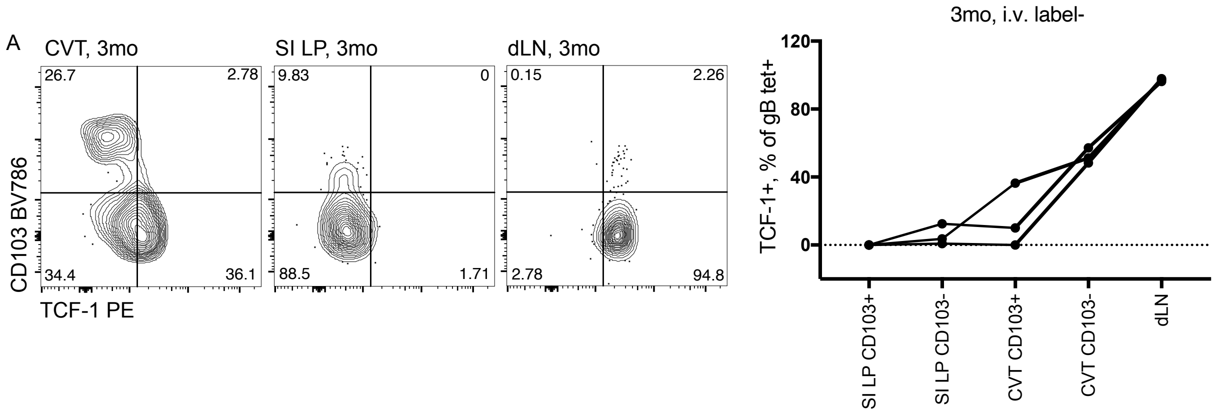


**Figure S2. Comparison of TCF-1 protein expression in HSV-specific CD8 T cells from mouse CVT and SI LP.** (A) Protein expression of CD103 and TCF-1 among HSV-specific i.v. label- CD8 T cells in CVT, SI LP, and dLN 3mo after LM-gB immunization. Flow plots represent concatenated data from 3 mice. (B) TCF-1+ frequency among CD103+ or CD103- subsets of HSV-specific CD8 T cells from CVT and SI LP compared to LN 3mo after immunization.

**Supplemental Figure 3**

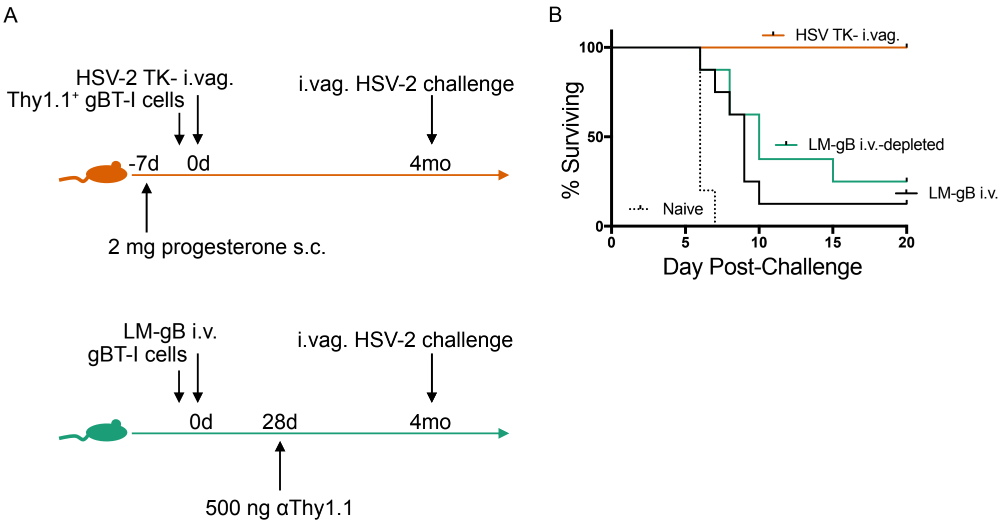


**Figure S3. Protection against HSV-2 challenge is not affected by circulating gBT-I depletion.** (A) Schematic of experiment. (B) Survival after vaginal HSV-2 challenge of naive and immunized mice.
